# Chronic potentiation of metabotropic glutamate receptor 2 with a nanobody accelerates amyloidogenesis in Alzheimer’s disease

**DOI:** 10.1101/2024.01.22.576777

**Authors:** Pierre-André Lafon, Mireille Elodie Tsitokana, Ugo Alenda, Clémentine Eva Philibert, Mathieu Oosterlaken, Marta Cimadevila, Jessica Monnic, Salomé Roux, Julie Bessié, Séverine Diem, Franck Vandermoere, Laurent Prézeau, Patrick Chames, Julie Kniazeff, Sylvie Claeysen, Jean-Philippe Pin, Véronique Perrier, Jianfeng Liu, Philippe Rondard

## Abstract

Immunotherapy of Alzheimer’s disease (AD) is a promising approach to reduce the accumulation of amyloid-beta (Aβ), a critical event in the onset of the disease. Targeting the group II metabotropic glutamate receptors, mGlu2 and mGlu3, could be important in controlling Aβ production, although their respective contribution remains unclear due to the lack of selective tools. Here, we show that enhancing mGlu2 receptor activity increases Aβ_1-42_ peptide production whereas activation of mGlu3 has no effect. We show that such a difference likely results from the direct interaction of APP with mGlu3, but not with mGlu2 receptors, that prevents APP amyloidogenic cleavage and Aβ_1-42_ peptides production. We then show that chronic treatments of the AD model 5xFAD mice with a brain-penetrating mGlu2-potentiating nanobody accelerated amyloid aggregation and exacerbated memory deficits, but had no effect in control mice. Our results confirm that a selective mGluR2 activation exacerbates AD disease development, suggesting that therapeutic benefices could be obtained with blockers of this receptor. Our study also provides the proof-of-concept that chronic administration of nanobodies targeting neuroreceptors can be envisioned to treat brain diseases.

## Introduction

Alzheimer’s disease (AD) is the most common neurodegenerative disorder characterized by progressive cognitive decline, memory impairments, and the accumulation of protein aggregates (amyloid plaques and Tau tangles)^1–3^. Amyloid deposits are mainly constituted of aggregated amyloid-β (Aβ) peptides which are considered the primary cause of this pathology^4^. Aβ peptides derive from the sequential cleavage of the amyloid precursor protein (APP) by the β-(BACE1) and γ-secretases^5^. These last decades, many small chemical drugs targeting AD’s markers have failed to demonstrate clinical efficacy^6^. Recently, passive immunotherapies based on Aducanumab and Lecanemab antibodies targeting Aβ peptides, were approved by the FDA and are used to treat AD patients with mild cognitive impairments and early dementia stages^7,8^. However, approval of Aducanumab was controversial^9^ as repeated injections and high doses led to amyloid-related imaging abnormalities (ARIA) in 43% of the treated-patients^10^. Thus, antibody-based therapy seems to be a promising strategy for improving AD, but new generations of antibodies with less secondary effects still need to be developed.

While extensive research helped to understand the underlying causes and mechanisms of AD, identifying new targets for the development of antibody-based therapies is urgently needed. Emerging evidences indicate that metabotropic glutamate receptors (mGluRs)^11^, belonging to G protein-coupled receptors (GPCRs) could be involved in the progression of AD, representing potential targets for new treatments^12–14^. Some studies suggest a potential role of the group II mGluRs (mGluR2 and mGluR3) in AD, although their role on Aβ production is not clear. Indeed, a selective activation of mGluR2 (LY566332) in cultured neurons increases Aβ toxicity while nonspecific activation of mGluR2 and mGluR3 (LY379268) rather provided neuroprotective effects^15^. Studies using isolated intact nerve terminals prepared from an AD mouse model, showed that non-selective activation of the group II mGluRs increases the production and secretion of Aβ_1-42_ peptides^16^. Due to the limited tools to specifically target mGluR2 or mGluR3, their precise contribution to AD and the resulting potential therapeutic implications needs to be clarified.

Camelid single-domain antibodies^17^ (also called V_HH_ or nanobodies) are new agents that can modulate the activity of receptors including GPCRs. Growing evidences show they can offer a novel and specific approach for the treatment of neurological diseases^18^, with reduced side effects and low immunogenicity^19^. We have recently developed nanobodies that selectively bind to various subtypes of mGluR, acting either as agonist or as a positive allosteric modulators (PAM)^20–22^. Furthermore, their small size (15 kDa) could facilitate their ability to pass the blood-brain barrier (BBB)^18^. We recently reported a nanobody, DN13-DN1, specific for mGluR2 and acting as a PAM^23^. Interestingly, DN13-DN1 crosses the BBB and can be detected in the brain up to 7 days post-injection, after a single injection of 10 mg/kg through intraperitoneal (i.p.) route. Thus, DN13-DN1 represented a potential treatment for schizophrenia as mGluR2 are involved in this pathology^24^. We showed that a single injection of DN13-DN1 rescues the cognitive deficits of a neonatal mouse model of schizophrenia induced with phencyclidine (PCP)^23^. Consequently, DN13-DN1 is a promising tool to determine the specific contribution of mGluR2 in the pathogenesis of AD.

In the present study, we show that activation of mGluR2, with a small drug or with the DN13-DN1 nanobody, increases the production and secretion of Aβ_1-42_ peptides in transfected cells. We demonstrated that mGluR2 is not interacting with APP, the latter being thus more prone to be internalized and cleaved by the β-secretase, enhancing Aβ levels. Conversely, mGluR3 interacts with APP, thus protecting it from β-cleavage and inhibiting Aβ_1-42_ production. Chronic treatment of an AD mouse model (5xFAD) through repeated i.p. injections of DN13-DN1, accelerates cognitive deficits associated with an increased amyloid load. Altogether, this study provides a proof-of-concept that targeting mGluR2 with a PAM nanobody can have measurable effects on both cognitive deficits and Aβ load. This also suggests that other mGluRs are potentially relevant therapeutic targets for AD and fine modulation of their function with nanobodies exhibiting allosteric properties could efficiently modify the course of AD.

## Results

### Activation of mGluR2 increases Aβ_1-42_ production, but not mGluR3

To decipher the potential roles of mGluR2 and mGluR3 on the production and secretion of Aβ_1-42_ peptides, we investigated the effect of their activation on APP processing in HEK293 cells and measured the Aβ_1-42_ by using a TR-FRET-based assay (**Figure 1A**). HEK293 cells, expressing an endogenous human APP (eAPP), were transfected with a snap-tagged mGluR2 or mGluR3 (^ST^mGluR2 or ^ST^mGluR3) constructs. ^ST^mGluR2 transfected cells treated with PBS as a control showed that Aβ_1-42_ peptides were not secreted in the cell supernatant (*p* > 0.05). However, activation of ^ST^mGluR2 increased significantly the production and secretion of Aβ_1-42_ in the supernatant compared to the control condition (Mock), when activated with the full agonist LY379268 (LY37, 5 µM, *p* < 0.0001), or with a nanobody specific to mGluR2 and exhibiting positive allosteric modulator (PAM) effects DN13-DN1 (200 nM, *p* < 0.0001), or with the EC_20_ of LY379268 (0.5 nM) alone (*p* = 0.0374) or combined with DN13-DN1 (*p* < 0.0001) (**Figure 1B**). To note, the DN13-DN1 nanobody alone increased the secretion of Aβ_1-42_ peptides due to the presence of ambient glutamate. Preincubation of cells with the antagonist LY341495 (LY34, 10 µM) before adding the agonist LY37 (5 µM) showed that inhibition of ^ST^mGluR2 significantly decreased the secreted Aβ_1-42_ levels (*p* < 0.0001) (**Figure 1B**). By contrast, treatment of cells transfected with ^ST^mGluR3 in a similar way as described above, did not induce any production and release of Aβ_1-42_ peptides (*p* > 0.05) (**Figure 1B**).

**Figure 1.**
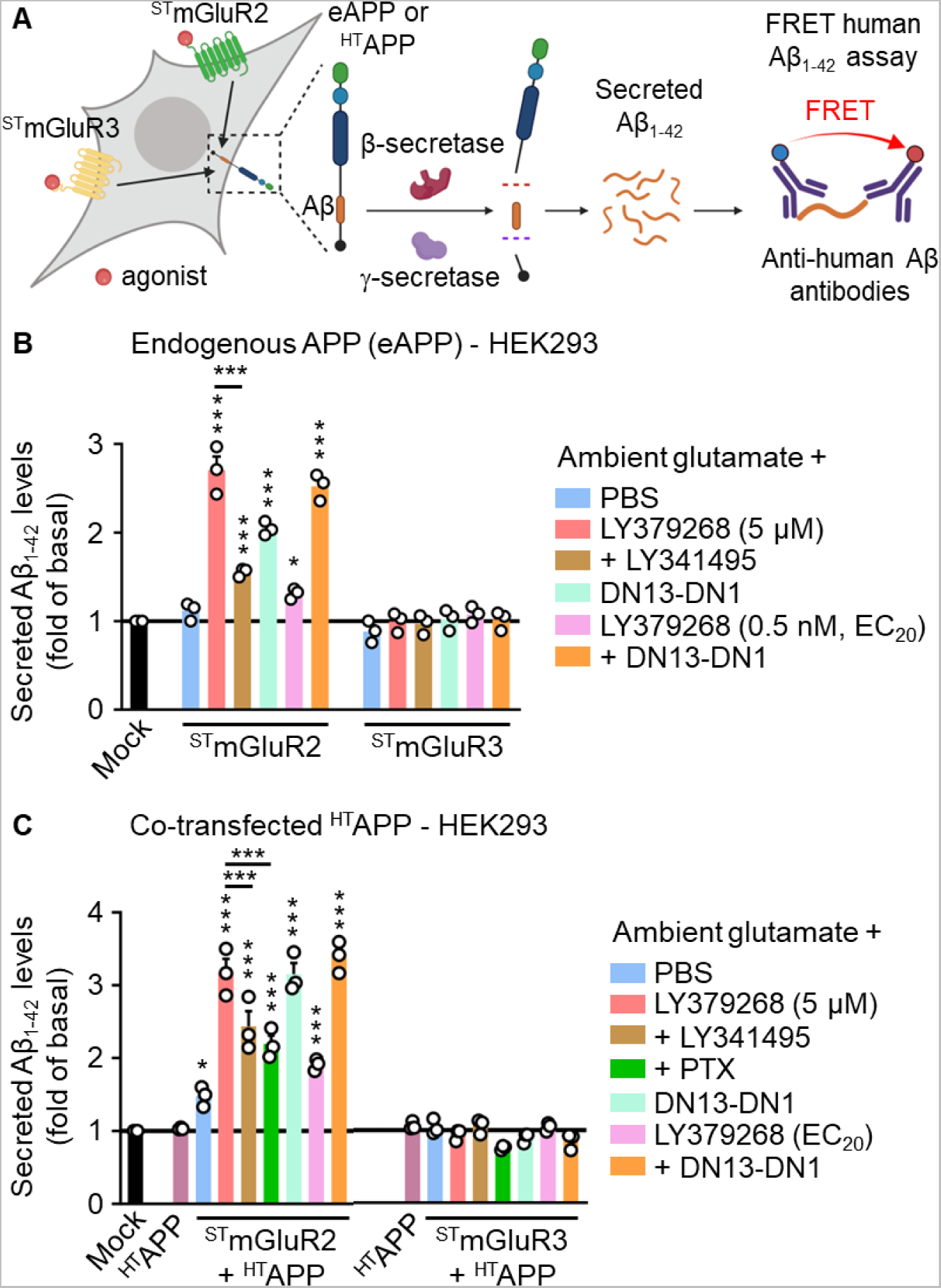
mGluR2 activation increases Aβ1-42 secretion in HEK293 cells. **A.** Schematic illustration of HEK293 cells transfected with human snap-tagged mGluR2 (^ST^mGluR2) or ^ST^mGluR3, in presence of the endogenous APP (eAPP) or co-transfected with human halo-tagged APP (^HT^APP). The impact of mGluR2 or mGluR3 activation on APP processing was performed by measuring secreted levels of Aβ_1-42_ in the conditioned medium through FRET-based assay. Created with BioRender.com. **B.** Secreted Aβ_1-42_ levels produced from the endogenous APP (eAPP) in HEK293 cells transfected with either empty vector (Mock), or ^ST^mGluR2, or ^ST^mGluR3. Cells were treated either with PBS; or with agonist LY379268 (5 µM) alone, or with a pre-treatment with the antagonist +LY341495 (10 µM); DN13-DN1 (200 nM); or with the EC_20_ of LY379268 (0.5 nM) alone, or combined with DN13-DN1. **C.** Secreted Aβ_1-42_ levels in HEK293 cells co-expressing ^HT^APP and either ^ST^mGluR2 or ^ST^mGluR3. Cells were treated either with PBS; LY379268 (5 µM) alone, or with a pre-treatment with LY341495 (10 µM), or with pertussis toxin (PTX, 0.2 µg/mL); DN13-DN1 (200 nM); or with the EC_20_ of LY379268 (0.5 nM) alone, or combined with DN13-DN1. As a control, cells were transfected with either the empty vector (Mock) or ^HT^APP. Data in **B-C** represent mean ± SEM from 3 biologically independent experiments performed in triplicates and are presented as a ratio of the basal levels. Data were analyzed using the ordinary one-way ANOVA followed by a Holm-Sidak’s *post hoc* analysis (* *p* < 0.05, *** *p* < 0.001).

Co-transfection of ^ST^mGluR2 with a human halo-tagged APP (^HT^APP) confirmed that activation of this receptor increases the production and secretion of Aβ_1-42_ peptides (*p* < 0.05), but not when ^HT^APP is co-transfected with ^ST^mGluR3 (*p* > 0.05) (**Figure 1C**). Activation of mGluR2 with LY37 (5 µM) induced a significant increase of secreted Aβ_1-42_ peptides compared to Mock condition (*p* < 0.0001). Interestingly, pre-incubation of cells either with LY34 (10 µM) or with the pertussis toxin (PTX, 0.2 µg/mL), inhibiting G_αi/o_ signaling, significantly reduced the secreted levels of Aβ_1-42_ upon activation of mGluR2 with LY37 (5 µM) (*p* < 0.0001) compared to LY37 (5 µM) condition alone (**Figure 1C**). The PTX pre-treatment suggests that activation of mGluR2 induces the production of Aβ_1-42_ peptides through a G-protein dependent pathway.

In parallel, we checked the cell surface expression of the different receptors transfected by using the fluorescent markers Snap- or Halo-Lumi4-Tb (**Sup. Figure 1A**). Expression was measured using the Tb emission at 620 nm, upon excitation at 337 nm. The cell surface expression of ^ST^mGluR2 and ^ST^mGluR3, transfected alone or co-transfected with ^HT^APP, was measured before adding any compound (t=0h) (**Sup. Figure 1B, D**) to determine the basal level of expression of these receptors. After 12h of incubation with the different compounds and nanobodies, cell surface expression of the different receptors showed that they were remaining expressed at the plasma membrane at sufficient levels (**Sup. Figure 1C, E-F**). This suggests that the absence of production of Aβ_1-42_ peptides in ^ST^mGluR3 transfected cells is not due to a decreased expression of this receptor upon activation.

Taken together these data indicate that activation of mGluR2 is implicated in the production and secretion of Aβ_1-42_ peptides mediated, in part, through a G-protein dependent manner. In contrast, mGluR3 does not induce amyloid peptide production upon activation.

### mGluR3 specifically associates with APP, but not mGluR2

We next examined why mGluR2 and mGluR3 act differently towards Aβ_1-42_ peptides’ production while sharing an amino acid sequence homology of approximately 70%^25^ and being coupled to the same G_α_ subunit. It has been described that APP can bind to the N-terminal sushi domain (SD1) of GB1a subunit of the GABA_B_ receptor and this complex decreases the availability of APP to be processed by the β-secretase, thus limiting the production of the Aβ peptides in the endosomal compartment^26^. As activation of mGluR3 does not produce Aβ_1-42_ peptides (**Figure 1B-C**), we hypothesized by analogy with the GABA_B_ receptor that mGluR3 may interact with APP, but not mGluR2. To assess the interaction between APP and mGluR2 or mGluR3, we performed FRET saturation assays in HEK293 cells (**Figure 2A**). In this assay, transfected Snap-tagged (ST) receptors were labelled with the donor (Snap-Lumi4-Tb) and Halo-tagged (HT) receptors with the acceptor (Halo-red). Under constant expression of ^ST^GB1a-ASA, the FRET signal increased hyperbolically as a function of ^HT^APP expression (black curve) (**Figure 2B and source data 1**), confirming the specific association of GB1a-APP described by Dinamarca and colleagues^26^. Cells co-expressing ^ST^APP and ^HT^mGluR3 (orange curve) exhibited a FRET saturation curve equivalent to the one observed for ^ST^mGluR2 with ^HT^mGluR2 (blue curve, positive control), subunits known to associate together ^27^, reflecting a specific interaction of mGluR3 with APP (**Figure 2B and source data 1**). In contrast, cells co-expressing ^ST^APP and ^HT^mGluR2 (red curve) exhibited a low FRET saturation curve, comparable to the one observed for cells expressing ^ST^mGluR2 with ^HT^mGluR5 (grey curve, negative control), two receptors that do not interact together to form heterodimers^27^. This collisional FRET shows that APP does not associate with mGluR2 (**Figure 2B and source data 1**).

**Figure 2.**
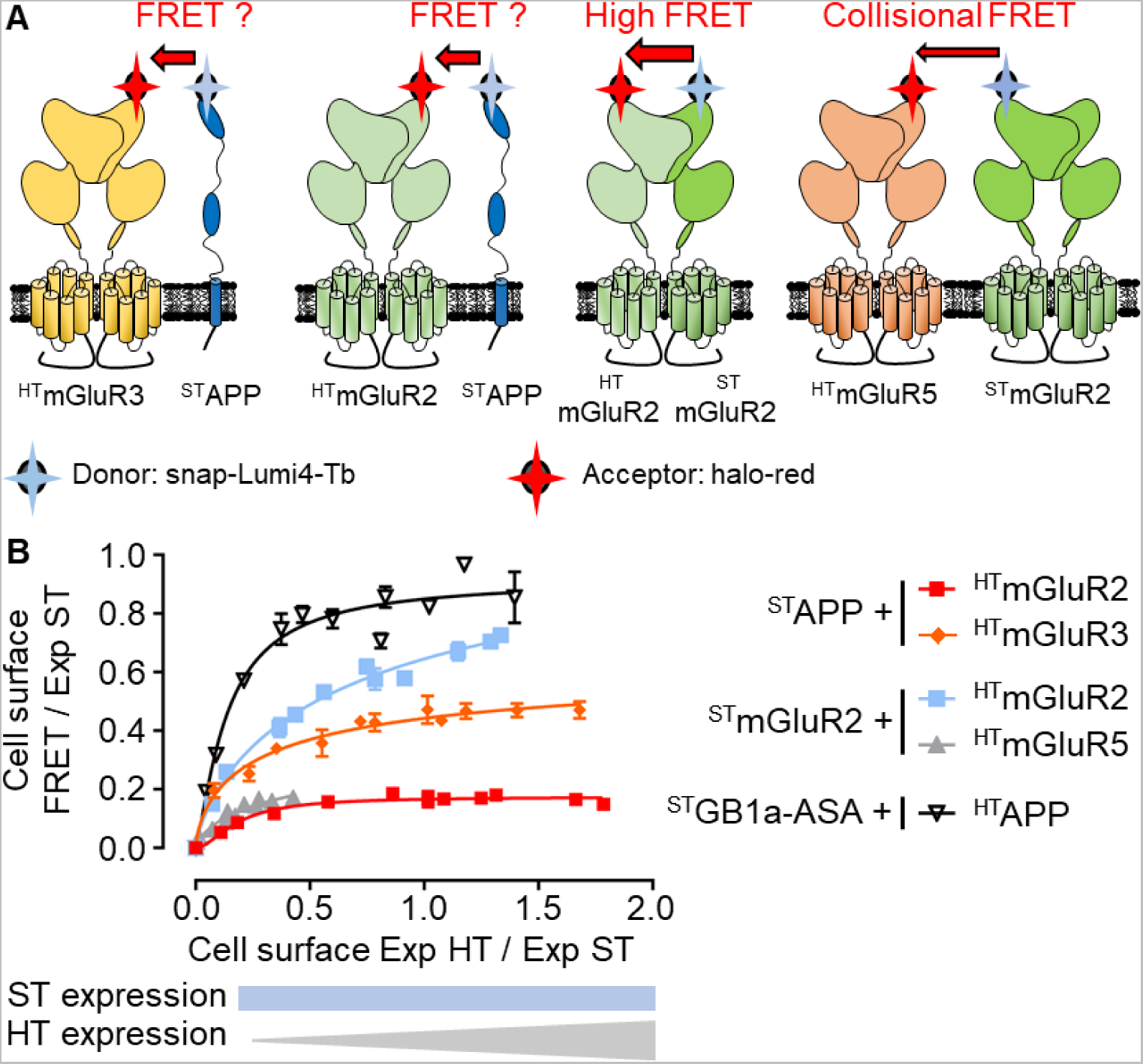
APP interacts with mGluR3 but not mGluR2. **A.** Schematic illustration of the TR-FRET saturation assay conditions used to determine if mGluR2 and mGluR3 can interact with APP. **B.** TR-FRET saturation curves in HEK293 cells of ^ST^APP with either ^HT^mGluR3 (orange) or ^HT^mGluR2 (red); ^ST^mGluR2 with ^HT^mGluR2 (blue, positive control) or with ^HT^mGluR5 (grey, negative control); and ^ST^GB1a-ASA with ^HT^APP (black, positive control). Snap-tagged constructs were transfected at a fix level and labelled with the donor (Snap-Lumi4-Tb), while Halo-tagged receptors were transfected with increasing quantities and labelled with the acceptor (Halo-red). Expression of cell surface snap-(Exp ST) and halo-tagged (Exp HT) receptors as well as the FRET signal were measured. FRET/Exp ST is shown as a function of the Halo to Snap expression ratio (Exp HT/Exp ST). Data are presented as mean ± SEM of triplicates from a representative experiment, performed 3 times.

To better characterize the APP-mGluR3 interaction, we performed FRET saturation assays with different APP and mGluR3 mutants. The FRET signal is highly decreased in cells co-expressing ^ST^APP and ^HT^mGluR3(Del570-879)-GPI (pink curve), corresponding to the deletion of the C-terminal and the transmembrane domains and inserted in the membrane with a GPI anchor. The FRET level with ^HT^mGluR3(Del570-879)-GPI mutant is comparable to the saturation obtained for ^ST^APP and ^HT^mGluR2 (red curve, negative control) (**Figure 3A and source data 2A**). This suggests that the C-terminal and the transmembrane domains of mGluR3 are involved in the interaction with APP. Indeed, a deletion of only the C-terminal domain of ^HT^mGluR3 (Del829-879) (green curve) decreased the FRET saturation of 50% compared to APP-mGluR3 (orange curve, positive control) but remains higher than the negative control (**Figure 3A and source data 2A**). In addition, deletion of the C-term domain of APP (Del648-695) co-transfected with ^ST^mGluR3 also shows a reduction of FRET signal (blue curve). These results suggest that the C-terminal domains of mGluR3 and APP are necessary for their interaction but are not sufficient.

**Figure 3.**
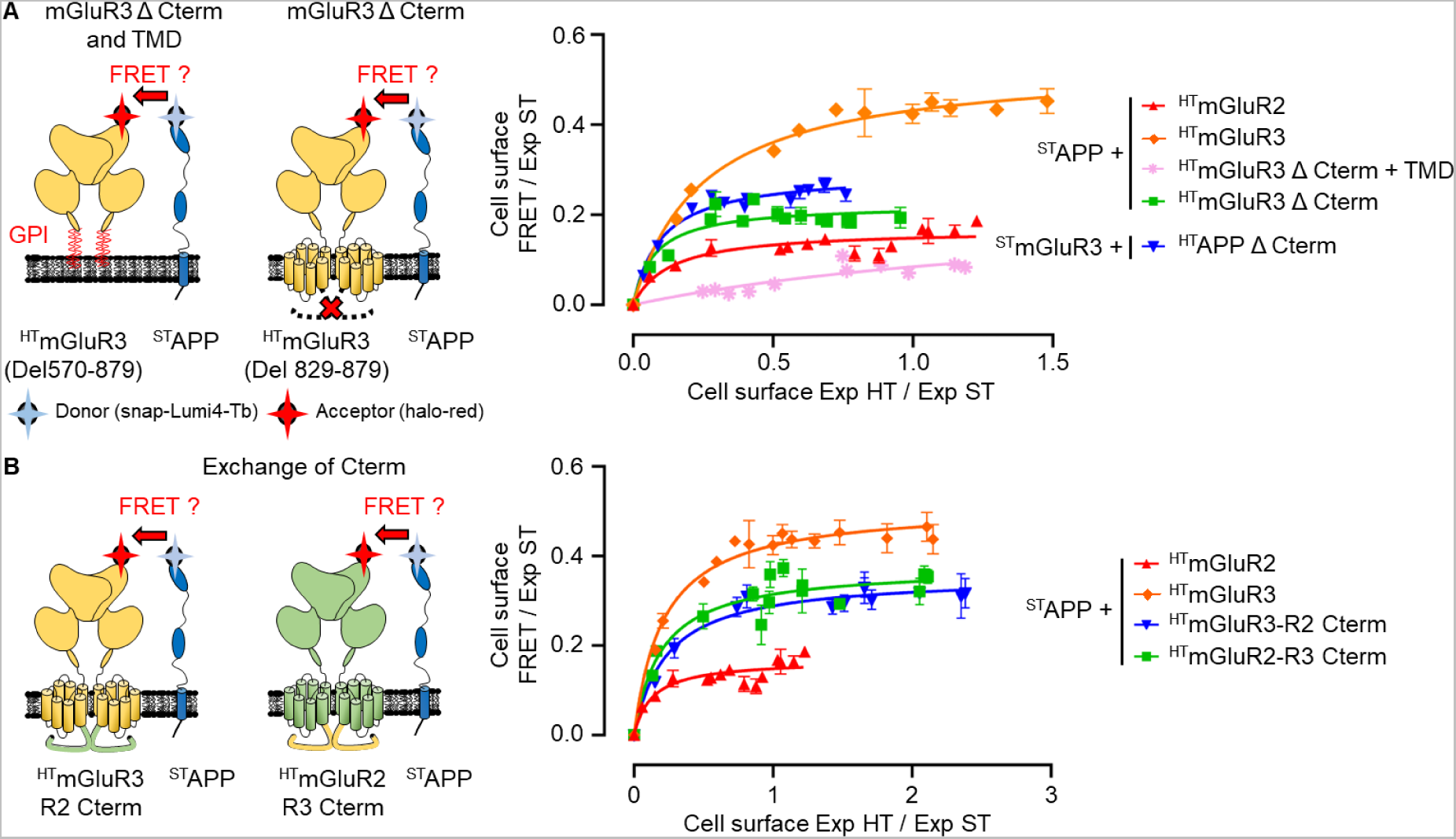
APP interacts with mGluR3 through the transmembrane and C-terminal domains. **A.** TR-FRET saturation curves in HEK293 cells of ^ST^APP with either ^HT^mGluR2 (red, negative control), or with ^HT^mGluR3 (orange, positive control); ^ST^APP with either ^HT^mGluR3(Del570-879) (pink), or ^HT^mGluR3(Del829-879) (green); and ^ST^mGluR3 with ^HT^APP(Del648-695) (blue). **B.** TR-FRET saturation curves in HEK293 cells of ^ST^APP with either ^HT^mGluR2 (red, negative control) or with ^HT^mGluR3 (orange, positive control); and ^ST^APP with either ^HT^mGluR3-R2 Cterm (blue) or ^HT^mGluR2-R3 Cterm (green). All Snap-tagged constructs were transfected at a fix level and labelled with the donor (Snap-Lumi4-Tb), while all Halo-tagged receptors were transfected with increasing quantities of cDNA and labelled with the acceptor (Halo-red). Expression of cell surface snap-(Exp ST) and halo-tagged (Exp HT) receptors, as well as the FRET signal were measured. FRET/Exp ST is shown as a function of the Halo to Snap expression ratio (Exp HT/Exp ST). Data are presented as mean ± SEM of triplicates from a representative experiment, performed 3 times.

Since the deletion of the C-terminal domain of mGluR3 could also destabilize its transmembrane domain, we performed FRET saturation assays by exchanging the C-terminal domains between mGluR2 and mGluR3 (**Figure 3B and source data 2B**). FRET signals of ^ST^APP with ^HT^mGluR2-R3 Cterm (green curve), or ^ST^APP with ^HT^mGluR3-R2 Cterm (blue curve) (**Figure 3B and source data 2B**), are comparable to those seen in mGluR3 (Del829-879) (green curve in **Figure 3A and source data 2A**), suggesting a potential role of the transmembrane domains of mGluR3 and APP in their association. Altogether, these data indicate that mGluR3 specifically associates with APP, but not with mGluR2, and suggest that this interaction is mediated both through the C-terminal and transmembrane domains.

### APP is internalized upon mGluR2 activation

We then performed an internalization assay using the diffusion-enhanced resonance energy transfer (DERET)^28^, on HEK293 cells expressing mGluR2 or mGluR3 to determine the internalization profile of APP upon activation of these receptors. To this purpose, ^HT^APP, alone or co-transfected with ^ST^mGluR2 or ^ST^mGluR3, were labeled with Snap- or Halo-Lumi4-Tb and incubated with an excess of fluorescein (**Sup. Figure 2A**). Fluorescein generates the quenching of the Tb emission when receptors are present at the plasma membrane. Upon internalization of the receptors, the Tb emission is recovered allowing the measurement of their internalization over time, as fluorescein does not cross the plasma membrane (**Sup. Figure 2A**). Before starting the experiments, cell surface expression of ^ST^mGluR2, ^ST^mGluR3 and ^HT^APP, alone or in co-transfection, was monitored and similar expression of these receptors was measured, prior to assess their internalization profiles (**Sup. Figure 2B-C**).

To assess the internalization of APP upon mGluR2 or mGluR3 activation, we performed DERET internalization assay as described above (**Figure 4**). At the basal state (PBS), co-transfection of ^ST^mGluR2 with ^HT^APP did not modify their constitutive internalization profile (dark blue curves) (**Figure 4A**), showing that mGluR2 does not influence APP internalization (**Figure 4B**). However, activation of ^ST^mGluR2 with LY37 (1 µM) increases ^HT^APP internalization (red curves), suggesting that APP is addressed to the endosomal compartment to be processed into Aβ peptides, upon mGluR2 activation (**Figure 4A-B, right panels**).

**Figure 4.**
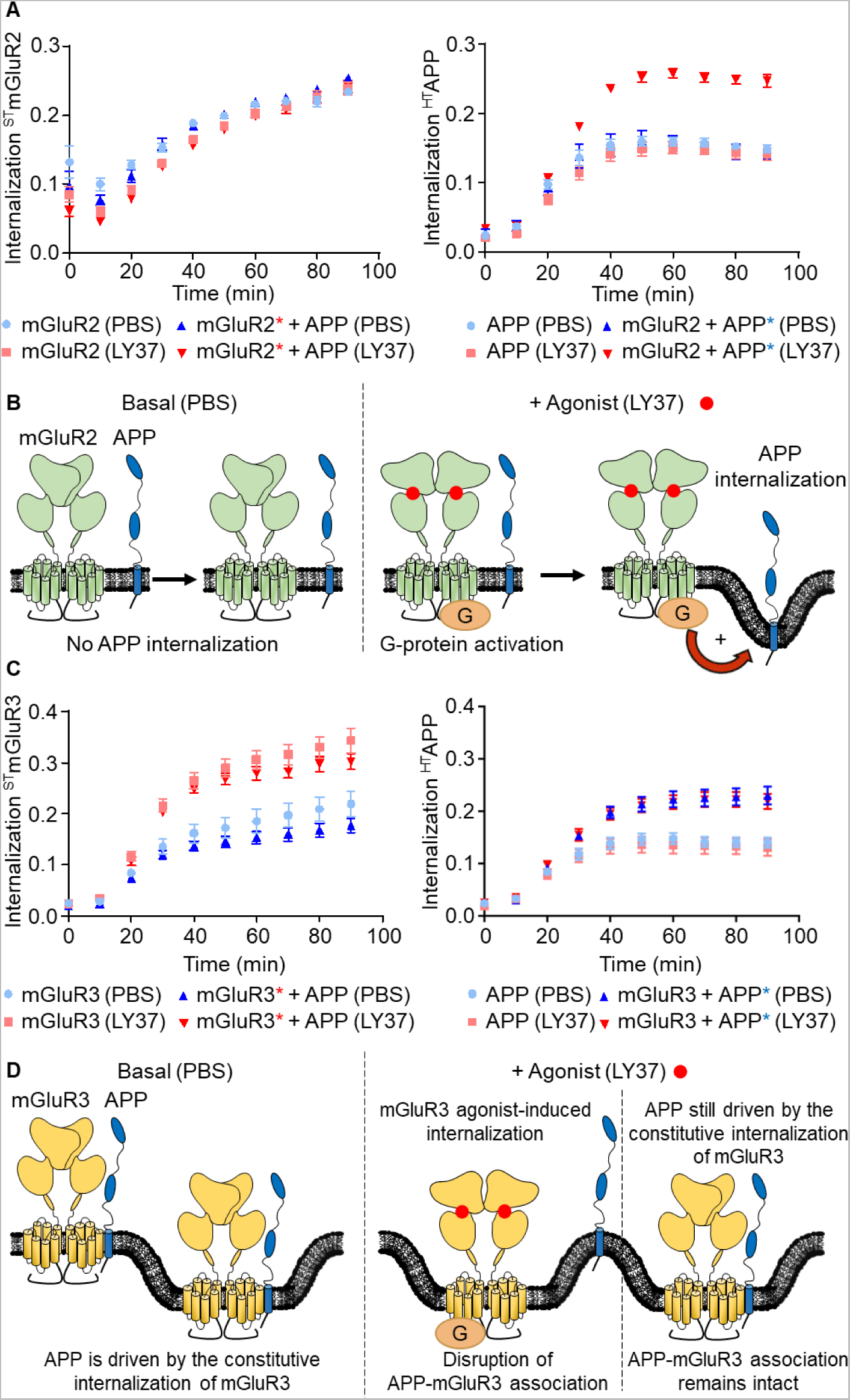
APP is internalized upon mGluR2 activation. **A.** ^ST^mGluR2 and ^HT^APP where transfected separately or together and treated with PBS (light and dark blue, respectively) or with 1 µM of the agonist LY379268 (pink and red, respectively). We measured by DERET the internalization of ^ST^mGluR2 (Left panel) and ^HT^APP (right panel). Data are presented as mean ± SEM from 3 biologically independent experiments performed in triplicates. **B.** At the basal state, mGluR2 does not modify APP internalization (left panel). When activated by an agonist (LY37), mGluR2 activation increases APP internalization (right panel), through a G protein dependent mechanism (Cf. also Figure 1). **C.** Internalization profiles of ^ST^mGluR3 and ^HT^APP transfected separately or together, treated with PBS (light and dark blue, respectively) or with 1 µM of the agonist LY379268 (pink and red, respectively). Data are presented as mean ± SEM from 3 biologically independent experiments performed in triplicates. **D.** At the basal state, in presence of mGluR3, APP is internalized since APP is driven by the constitutive internalization of mGluR3 due to the APP-mGluR3 association. When mGluR3 is activated, the complex APP-mGluR3 is disrupted and mGluR3 alone is agonist-induced internalized without APP. However, a part of the APP-mGluR3 complexes remains intact, not disrupted, and APP is still driven by the constitutive internalization of mGluR3.

Co-transfection of ^ST^mGluR3 with ^HT^APP did not modify mGluR3 constitutive internalization profile at the basal state (dark blue curve) nor its agonist-induced internalization capacity upon addition of LY37 (red curve) (**Figure 4C, left panel**). However, in presence of ^ST^mGluR3, ^HT^APP is driven by the constitutive internalization of mGluR3 at the basal state due to their interaction (**Figure 4D, left panel**) as shown by the increase of FRET (dark blue curve) (**Figure 4C, right panel**). Surprisingly, addition of the mGluR3 agonist LY37, did not further increased APP internalization (red curve) (**Figure 4C, right panel**) when both receptors are co-expressed. This suggests that when mGluR3 is activated, the complex APP-mGluR3 is disrupted and only mGluR3 undergoes agonist-induced internalization (**Figure 4D, right panel**). However, APP internalization profile co-transfected with mGluR3 was similar either at the basal state (PBS) or after addition of LY37 (dark blue and red curves) (**Figure 4C, right panel**). This result suggests that after addition of agonist, part of the APP-mGluR3 complexes remains intact and APP is still driven by the constitutive internalization of a subpopulation of mGluR3 (**Figure 4D, right panel**).

The combined data of these assays showed that APP internalization profile can be modified by mGluR2 and mGluR3. Activation of mGluR2 increases APP internalization suggesting an increased availability of APP to be processed into Aβ peptides in the endosomal compartment. In contrast, APP internalization driven by the constitutive internalization of mGluR3 rather suggests that the complex APP-mGluR3 protects APP from the β-cleavage after internalization.

### Chronic administration of DN13-DN1 nanobodies aggravates cognitive deficits in 5xFAD mice

Previously, we optimized a nanobody specifically targeting mGluR2 (DN13-DN1) that exhibits PAM properties *in vitro* and *in vivo*^23^. DN13-DN1 can cross the BBB after a single peripheral injection and is detected in the brain up to 7 days post-injection^23^. To determine the effects of a chronic positive modulation of mGluR2 in an AD context, we performed a long-term treatment of WT and 5xFAD male mice with DN13-DN1 at 10 mg/kg by i.p. injections once a week, for 22 weeks (**Figure 5A**). As negative controls, mice were treated for the same duration, either with 10 mg/kg of DN1, a nanobody specific of mGluR2 without any pharmacological properties^22^, or with an equivalent volume of PBS. A follow-up of WT and 5xFAD mice did not reveal any weight modification during the chronic treatment with the nanobodies when compared to their vehicle-treated littermates (**Sup. Figure 3A**). However, chronic administration of DN13-DN1 and DN1 nanobodies induced about 20% mortality for both WT and 5xFAD mice between 4 and 8 weeks of treatment (**Sup. Figure 3B**). A toxicological study performed on 27 organs collected in WT mice chronically treated with DN13-DN1, DN1 or with vehicle, did not induce any apparent toxicity of any organs examined. We observed some tubular basophilia in the kidneys of DN13-DN1 and DN1 treated animals, without any other concomitant changes indicative of toxicity. Its origin may be related to alterations in functional activity of the tubular cells rather than toxicity (data not shown).

**Figure 5.**
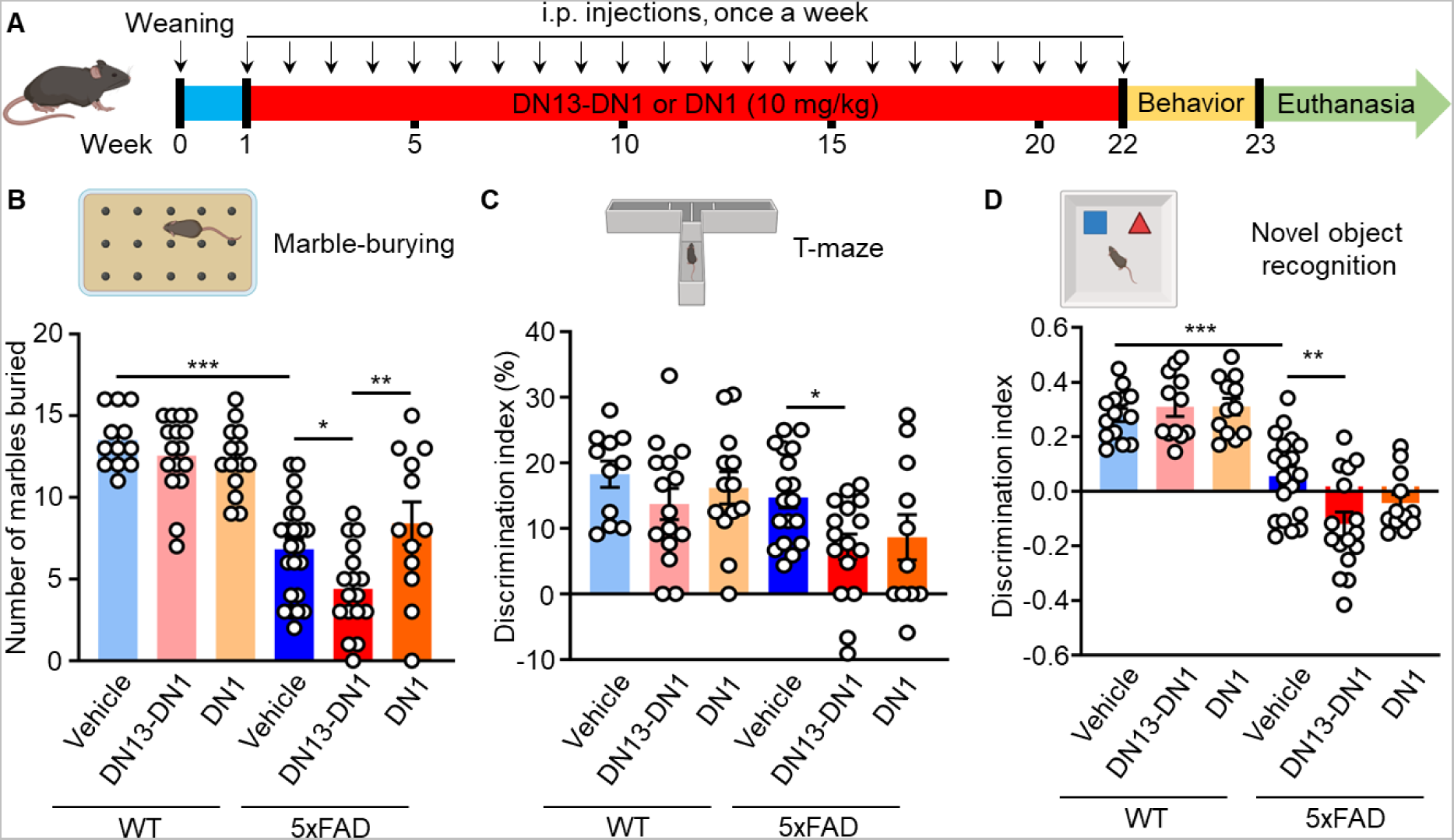
Positive modulation of mGluR2 using DN13-DN1 nanobody aggravates cognitive deficits in 5xFAD mice. **A.** Study design of the treatment of 5xFAD mice with nanobodies. One week after weaning, 5xFAD and WT male mice were chronically treated with 10 mg/kg of either mGluR2 positive allosteric modulator DN13-DN1 nanobody, or mGluR2 silent allosteric ligand DN1 nanobody, or with an equivalent volume of vehicle. Mice were i.p. injected once a week for 22 weeks. At the end of the treatment, mice underwent several behavioral tests such as nest-building, marble-burying, T-maze, and the novel object recognition. Created with BioRender.com. **B-D.** Number of marbles buried in the marble burying test (**B**), discrimination index in the T-maze test (**C**) and discrimination index in the novel object recognition test (**D**) were assessed for WT and 5xFAD mice treated with vehicle, 10 mg/kg of DN13-DN1 or with 10 mg/kg of DN1 nanobodies. Data are presented as mean ± SEM (n = 12-21 mice / group) and were analyzed using the two-way ANOVA followed by a Holm-Sidak’s *post hoc* analysis (\**p* < 0.05, \*\**p* < 0.01, \*\*\**p* < 0.001).

To determine whether chronic positive modulation of mGluR2 influences behavioral deficits in 5xFAD mice, we performed different behavioral tests. We first assessed the ability of animals to build nests^29^, a natural behavior of mice that requires hippocampal neuronal functions and reports on hippocampal integrity^30^. At 6.5 months of age, analyses indicated a significant effect of the genotype, in which the 5xFAD vehicle-treated mice exhibited lower scores in nest-building (*p* = 0.0384) (**Sup. Figure 4A**) and the quantity of shredded nestlet (*p* = 0.004) (**Sup. Figure 4B**) compared to WT mice treated with vehicle. However, chronic treatment of 5xFAD male mice with the DN13-DN1 or DN1 nanobodies did not further impact the nest quality (*p* > 0.05) (**Sup. Figure 4A-B**). This suggests that the chronic modulation of mGluR2 is not affecting, directly or indirectly, the hippocampal subpopulation of neurons responsible of the nest-building behavior in 5xFAD mice.

To measure the apathy-like behavior we used the marble-burying test^31^. Analyses revealed that 5xFAD vehicle-treated mice significantly buried fewer marbles than their WT littermates (*p* < 0.0001) and this phenotype is exacerbated when 5xFAD were treated with DN13-DN1 (*p* = 0.0472) (**Figure 5B**). This suggests that chronic treatment with a PAM mGluR2 aggravates the apathy-like behavior in 5xFAD mice. In contrast, DN1 treatment of 5xFAD mice did not change the number of buried marbles compared to 5xFAD mice treated with vehicle (*p* > 0.05) (**Figure 5B**). To note, treatment of WT mice with the nanobodies did not modify the burying behavior (*p* > 0.05) (**Figure 5 B**), suggesting that chronic positive modulation of mGluR2 only exacerbate apathy-like behavior in a disease context.

We then evaluated the spatial and short-term memories of the animals at 6.5 months, by using the T-maze test, that involves the hippocampus^32,33^, crucial for spatial navigation and memory. No modification of the discrimination index was observed between WT and 5xFAD mice treated with vehicle (*p* = 9065) (**Figure 5C**). However, chronic treatment of transgenic mice with DN13-DN1 significantly decreased the discrimination index, reflecting fewer entries in the novel arm, compared 5xFAD vehicle mice (*p* = 0.049) (**Figure 5C**), suggesting that DN13-DN1 accelerates spatial and short-term memory deficits in 5xFAD mice. In addition, no effect of the chronic treatments with nanobodies was observed for WT mice (*p* > 0.05) on spatial and short-term memory (**Figure 5C**).

Finally, we assessed the long-term memory with the novel object recognition test, completed over 3 days: habituation day (empty arena), training day (2 identical objects), and testing day (1 novel object introduced). During the habituation phase, mice were allowed to explore the empty arena for 5 min and several parameters were analyzed such as the velocity, the time spent in the center of the arena, the distance, and the immobility. For all these parameters analyzed, except for the time spent in the center of the arena (*p* = 0.9763) (**Sup. Figure 4C**), we observed a main effect of the genotype in which 5xFAD vehicle-treated mice had a lower mean velocity (*p* = 0.0491) (**Sup. Figure 4D**), travelled lower distances in the arena (*p* = 0.0499) (**Sup. Figure 4E**) and exhibited an increased time of immobility (%) (*p* = 0.0438) (**Sup. Figure 4F**), compared to their WT littermates treated with vehicle.

During the testing day, one familiar object was replaced by a novel one 24 h after the familiarization, as expected we observed a genotype effect for the discrimination index where 5xFAD vehicle-treated mice explore less the novel object compared to their WT littermates treated with vehicle (*p* = 0.0009) (**Figure 5D**). However, this phenotype was exacerbated for the 5xFAD mice chronically treated with 10 mg/kg of DN13-DN1 (*p* = 0.0048) (**Figure 5D**) compared to the 5xFAD vehicle-treated mice. This suggests that chronic positive modulation of mGluR2 in 5xFAD mice accelerates the long-term memory deficits. Altogether, these data show that the chronic positive modulation of mGluR2 in an AD context, aggravates the apathy-like behavior, the spatial memory, and the short- and long-term memories.

### Chronic modulation of mGluR2 with the DN13-DN1 nanobody increases amyloid load in 5xFAD mice

We next examined the effects of the chronic positive modulation of mGluR2 with the DN13-DN1 nanobody on the amyloid load in the WT and 5xFAD animals (**Figure 6**). We first analyzed the aggregates on hemibrain tissue sections using the Thioflavin T (ThT) staining (**Sup. Figure 5**). We observed that chronic injections of DN13-DN1 in 5xFAD substantially increased the number of amyloid plaques (+ 52%, *p* = 0.0023) (**Figure 6A-B**) as well as the surface of the aggregates (+ 49%, *p* = 0.0136) (**Figure 6A, C**) in the hippocampus compared to 5xFAD vehicle-treated mice. No impact on the amyloid aggregates was observed upon chronic treatment of 5xFAD mice with the DN1 nanobody (*p* > 0.99) nor any impact on the WT mice chronically treated with DN13-DN1 or DN1 nanobodies (*p* > 0.99). Similarly, DN13-DN1 treatment in 5xFAD mice induced, in the cortex (**Figure 6D**), a significant increase of the number of aggregates (+36%, *p* = 0.0372) (**Figure 6E**) but not their surface (*p* = 0.5108) (**Figure 6F**) compared to 5xFAD mice treated with vehicle. As for the hippocampus, no impact on the cortical amyloid aggregates was observed upon chronic treatment of 5xFAD mice with the DN1 nanobody (*p* > 0.99) nor any impact on the WT mice chronically treated with the 2 nanobodies (*p* > 0.99) (**Figure 6E-F**). In addition, 5xFAD mice treated with DN13-DN1 exhibited a high intraneuronal ThT fluorescence in cortical neurons compared to 5xFAD mice treated with vehicle (**Figure 6D, enlargements in lower panel**). Similarly, DN1-treated transgenic mice also exhibited intraneuronal ThT fluorescence but in a lower extent compared to mice treated with DN13-DN1. These data suggest that the chronic positive modulation of mGluR2 through the DN13-DN1 nanobody may impact the production of Aβ peptides leading to increased aggregation processes in 5xFAD mice.

**Figure 6.**
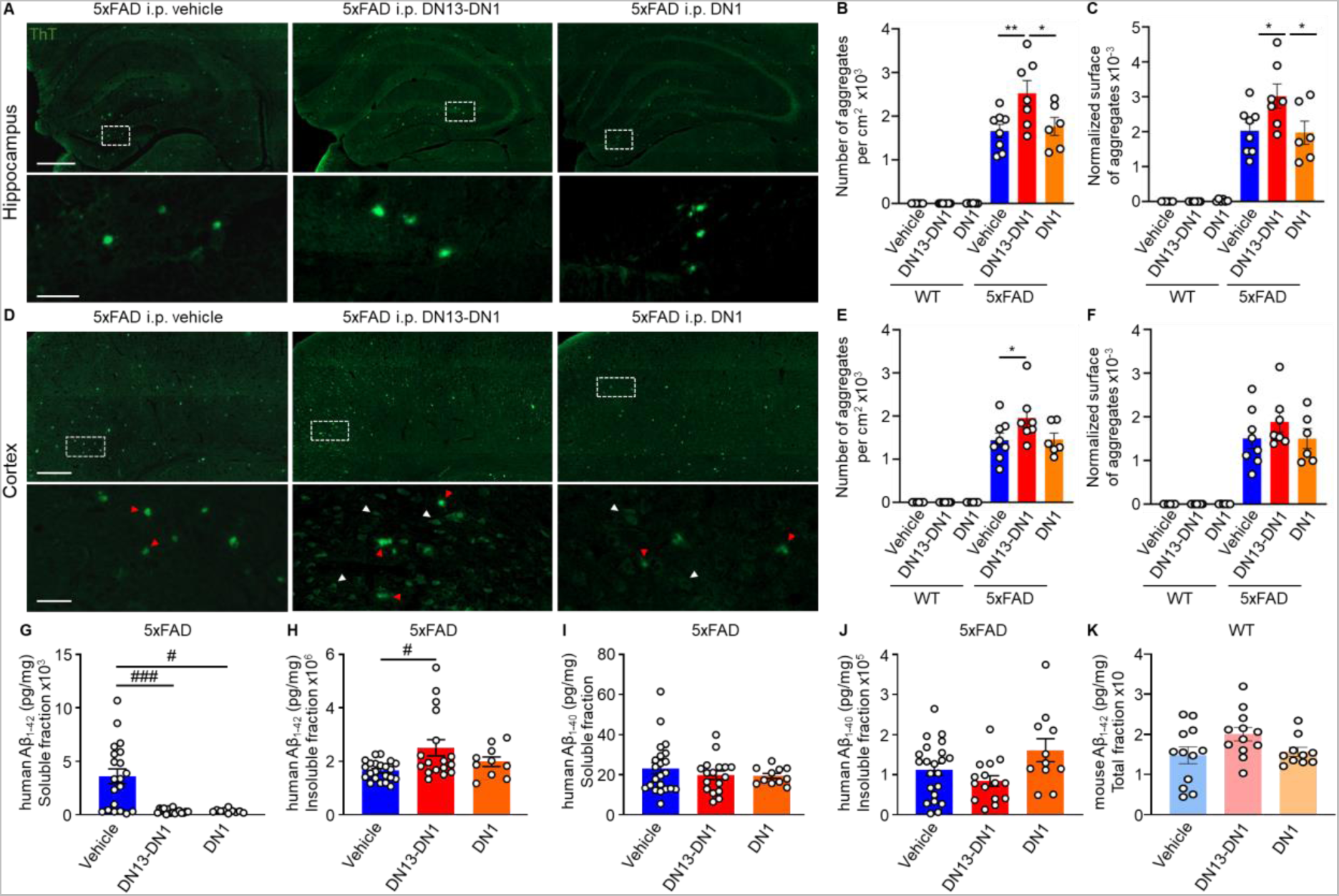
DN13-DN1 nanobody treatment worsen amyloid burden in 5xFAD mice. **A.** Representative images of thioflavin T (ThT) staining of amyloid plaques in the hippocampus of 5xFAD mice (27 weeks old) chronically treated with 10 mg/kg of either DN13-DN1, or DN1 or with an equivalent volume of vehicle. Images are mosaics obtained using a slide scanner Axio scan Z1 microscope with a 20X objective (scale bars: 200 and 50 µm). **B-C.** Quantification of the number (**B**) and the surface of aggregates (**C**) normalized by the total surface analyzed, in the hippocampus of WT and 5xFAD mice. Data are presented as mean ± SEM (n = 6-8 animals/group, 13-23 sections/animal) and analyzed using the two-way ANOVA followed by a Holm-Sidak’s *post hoc* analysis (\**p* < 0.05, \*\**p* < 0.01). **D.** Representative images of positive ThT amyloid plaques in the cortex of 5xFAD mice (27 weeks old) chronically treated with 10 mg/kg of DN13-DN1, 10 mg/kg of DN1 or with vehicle. Images are mosaics obtained using a slide scanner Axio scan Z1 microscope with a 20X objective (scale bars: 200, 50 and 10 µm). **E-F.** Quantification of the number (**E**) and the surface of aggregates (**F**) normalized by the total surface of cortex analyzed in WT and 5xFAD mice. Data are presented as mean ± SEM (n = 6-8 animals/group, 13-23 sections/animal) and analyzed using the two-way ANOVA followed by a Holm-Sidak’s *post hoc* analysis (\**p* < 0.05). **G-J.** Quantification of the human Aβ_1-42_ and Aβ_1-40_ levels in the soluble (**G, I**) and guanidine-treated insoluble fractions (**H, J**) from 5xFAD hemibrains, using a FRET-based assay. Data are presented as mean ± SEM (n = 11-22 animals/group) and analyzed using Kruskal-Wallis followed by a Dunn’s multiple comparison (**G-I**, #*p* < 0.05, ##*p* < 0.01) or the one-way ANOVA followed by a Holm-Sidak’s *post hoc* analysis (**J**, *p*> 0.05). **K.** Quantification of the total endogenous mouse Aβ_1-42_ levels from WT hemibrains, using an Elisa assay. Data are presented as mean ± SEM (n = 10-12 animals/group) and analyzed using Kruskal-Wallis followed by a Dunn’s multiple comparison.

Thus, we also assayed Aβ_1-42_ and Aβ_1-40_ levels in total brain homogenates of WT and 5xFAD mice. Surprisingly, 5xFAD mice chronically treated with DN13-DN1 and DN1 nanobodies exhibited a strong reduction of Aβ_1-42_ soluble fraction on brain homogenates compared to control animals (*p* = 0.0008 and *p* = 0.0178, respectively) (**Figure 6G**). Consistent with the increased amyloid plaque load, quantification showed a clear increase of the insoluble Aβ_1-42_ fraction in brain samples from 5xFAD mice treated with the DN13-DN1 nanobody (+52%, *p* = 0.0475) compared to 5xFAD control mice (**Figure 6H**). DN1 nanobody treatment did not modify the Aβ_1-42_ insoluble levels compared to vehicle-treated mice (*p* = 0.3908) (**Figure 6H**). Regarding Aβ_1-40_ levels on total brain homogenates no differences were observed upon chronic administration of DN13-DN1 or DN1 nanobodies for both soluble (*p* > 0.99) (**Figure 6I**) and insoluble (*p* = 0.2366 and *p* = 0.1464, respectively) (**Figure 6J**) fractions, compared to 5xFAD control mice. Interestingly, chronic treatment of WT mice with the nanobodies did not modify the total mouse Aβ_1-42_ levels compared to the WT vehicle-treated mice (*p* = 0.1780 for DN13-DN1 and *p* > 0.99 for DN1) (**Figure 6K**). Altogether, these findings suggest that DN13-DN1 chronic administration impacts the total brain Aβ_1-42_ content in 5xFAD mice.

### DN13-DN1 nanobody treatment decreases the relative expression of the mGluR2 homodimers in the cortex of 5xFAD mice

DN13-DN1 treatment had a lower impact on the amyloid load in the cortex of 5xFAD mice compared to the hippocampus, thus we examined whether the expression of mGluR2 could have been related to this difference (**Figure 7**). To this purpose, we quantified the relative expression of mGluR2 homodimers at the cell membrane by using DN1-Tb and DN1-d2 nanobodies in different brain regions using FRET technology (**Figure 7A**). As a negative control, we used the midbrain as mGluR2 is not expressed in this region^20^ (**Figure 7B**). In the cortex, we observed that 5xFAD mice treated with vehicle have less mGluR2 homodimers compared to WT animals (*p* = 0.0129) (**Figure 7C**), which is exacerbated upon DN13-DN1 treatment (*p* = 0.0014) (**Figure 7C**). No modifications of mGluR2 expression were recorded in the hippocampus (*p* > 0.05) (**Figure 7D**) or the cerebellum (*p* > 0.05) (**Figure 7E**) between 5xFAD-treated and WT mice. To note, chronic treatment with the nanobodies did not modify the expression of mGluR2 homodimers in WT mice (**Figure 7A-E**). The representative slopes calculated for mGluR2 relative quantification in the different conditions are presented in **Sup. Figure 6**. Altogether, chronic administration of DN13-DN1 nanobodies in 5xFAD mice decreases the relative expression of mGluR2 homodimers in the cortex only.

**Figure 7.**
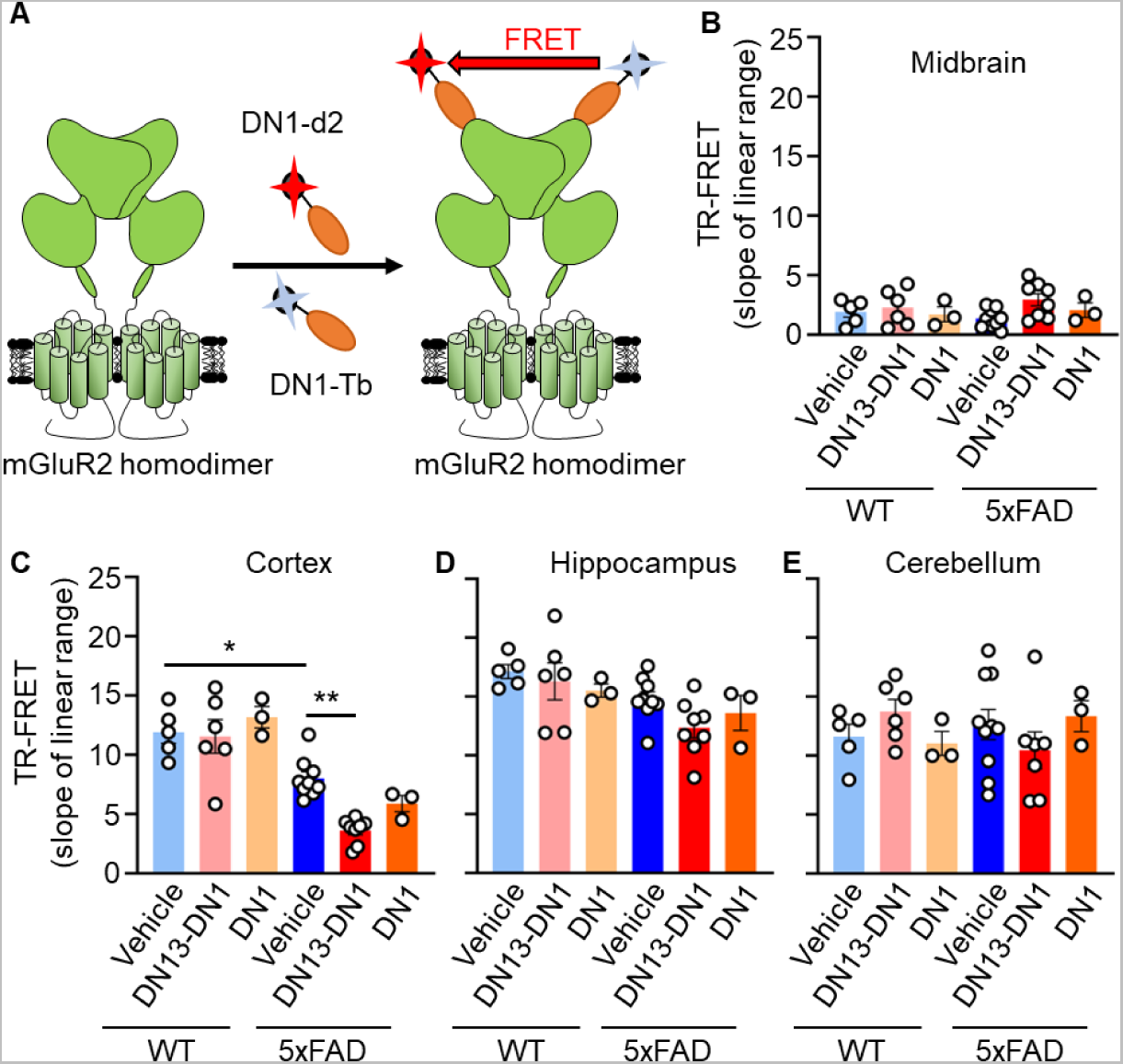
Reduction of mGluR2 homodimers in the cortex of 5xFAD mice upon DN13-DN1 nanobody treatment. **A.** Schematic representation of mGluR2 homodimers relative quantification with a FRET-based assay using DN1 nanobodies coupled to a donor (DN1-Tb) and acceptor (DN1-d2), both at 25 nM. **B-E.** Relative quantification of mGluR2 homodimers in the midbrain (**B**), cortex (**C**), hippocampus (**D**) and cerebellum (**E**) of WT and 5xFAD mice chronically treated with 10 mg/kg of either DN13-DN1 or DN1, or with vehicle. The TR-FRET signal indicates the slope values of the relative linear quantification experiments. Data are presented as mean ± SEM (n = 3-9 animals/group) and analyzed using the two-way ANOVA followed by a Holm-Sidak’s *post hoc* analysis (\**p* < 0.05, \*\**p* < 0.01).

## Discussion

AD is associated with an imbalance in glutamate signaling leading to excitotoxicity and neuronal damages^34^. Due to their location in the presynaptic compartment and roles in inhibiting glutamate release (G_αi/o_ signaling), activation of both mGluR2 and mGluR3 was believed to be neuroprotective against glutamate excitotoxicity in AD^35,36^. However, studies have demonstrated that non-selective activation of the group II mGluRs increases the production and secretion of Aβ_1-42_ peptides, but not of Aβ_1-40_, in isolated intact nerve terminals prepared from an AD mouse model^16^. This rather suggested that mGluR2 and/or mGluR3 can promote amyloidogenesis at the synapses, but their specific contribution was not assessed. In the present study, we brought clear evidence that mGluR2 is involved in the production of Aβ_1-42_ peptides through a G protein-dependent manner but not mGluR3. Indeed, mGluR3 has rather a protective role by interacting with APP through the transmembrane and C-terminal domains, decreasing the availability of APP to be processed into Aβ peptides in the endosomal compartments.

Since mGluR2 do not interact with APP (**Figure 2B**), receptor and G protein-activation induces the APP internalization (**Figure 4A-B**) and its processing by the β-secretase in the endosomal compartment. The non-association of mGluR2 with APP was also shown recently by a mGluR2 interactome of mouse prefrontal cortex using a specific mGluR2 nanobody and mass spectrometry analyses^37^. Interestingly, the production of Aβ_1-42_ peptides upon mGluR2 activation can have a physiological role. Indeed, physiological levels of Aβ_1-42_ peptides enhance learning and memory by favoring the long-term potentiation (LTP)^38–43^ (**Figure 8B**). This suggests that one possible source of production of physiological Aβ_1-42_ at the synapses comes from the activation of mGluR2 during strong and prolonged glutamate release, a prerequisite to activate the NMDA receptor-dependent LTP^34^. It is now well established that elevated Aβ peptide levels block the neuronal glutamate uptake leading to increased glutamate levels at the synaptic cleft^44,45^. Thus, in an AD context in which glutamate is released in excess, mGluR2 activation increases the production of Aβ_1-42_ peptides, which in turn blocks glutamate uptake perpetuating a vicious circle (**Figure 8A**).

**Figure 8.**
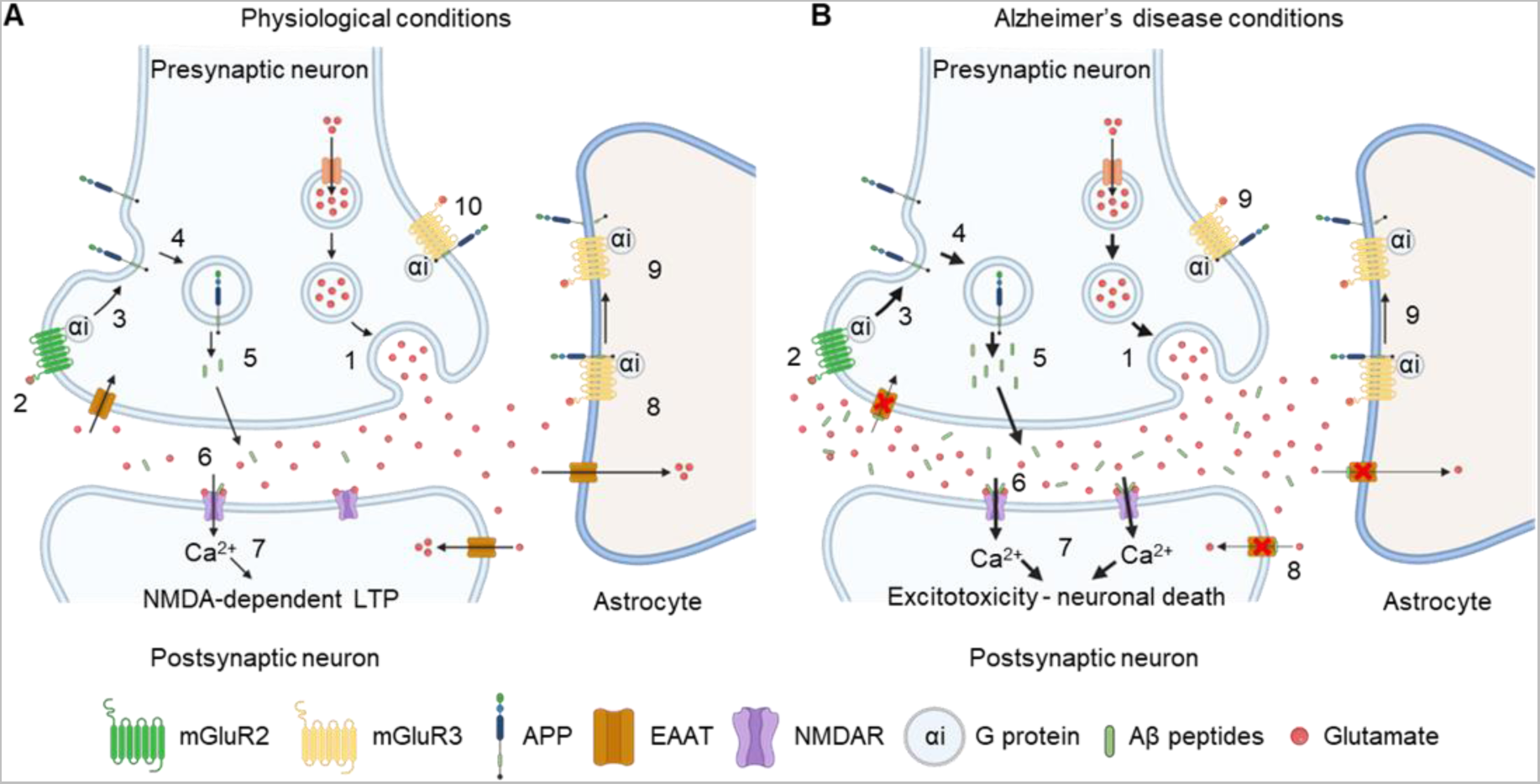
Proposed mechanisms for mGluR2 and mGluR3 actions towards APP processing. **A.** In physiological conditions, activation of the presynaptic neuron leads a glutamate release in synaptic cleft (**1**). Glutamate binds to mGluR2 (**2**) and its activation will lead to the internalization of APP (**3**). In the endosomal compartment (**4**), APP is cleaved by the β-secretase and generates Aβ peptides (**5**). These physiological levels of Aβ potentiate NMDA receptors at the synapse (**6**) favoring the NMDA-dependent long-term potentiation (LTP) (**7**). mGluR3 interacts with APP decreasing its availability to be processed by the β-secretase in the endosomal compartment. In astrocytes, activation of mGluR3 (**8**) promotes the non-amyloidogenic cleavage of APP by α-secretases resulting in the inhibition of Aβ peptides production (**9**). We hypothesize that neuronal mGluR3 behaves the same way as astroglial mGluR3 (**10**). **B.** In pathological conditions such as AD, glutamate is released in excess (**1**) and leads to strong and prolonged activation of mGluR2 (**2, 3**). This process will increase APP internalization (**4**) and β-processing, thus increasing Aβ levels at the synapse (**5**). With the excess of glutamate, Aβ peptides will reinforce the potentiation of NMDA receptors (**6**) leading to excitotoxicity that may cause neuronal death (**7**). Aβ peptides also contribute to the increased levels of glutamate at the synapse by blocking EAAT transporters, inhibiting glutamate recycling (**8**). For mGluR3, we hypothesize that its activation still enhances the α-processing of APP (**9**) as very limited studies investigated this mGluR3 mechanism in AD conditions.

By contrast, activation of mGluR3 does not induce the production of Aβ_1-42_ peptides (**Figure 1B-C**) due to its capacity to interact with APP (**Figure 2B**) through the C-terminal and transmembrane domains (**Figure 3**). This binding suggests an increased resident time of APP at the plasma membrane and a decreased availability of APP for β-secretase cleavage. Our data shows that internalization of APP can be driven by the constitutive internalization of mGluR3 (**Figure 4C-D**), due to their interaction, but this process does not induce the β-processing of APP (**Figure 8A**). This suggest that mGluR3-APP interaction induces a sterical hindrance preventing the β-secretase binding and cleavage of APP into Aβ peptides. We showed that in presence of agonist, mGluR3 undergoes agonist-induced internalization alone illustrating a disruption of the complex mGluR3-APP and suggesting that APP remains at the plasma membrane (**Figure 4C-D**). Previous studies showed that activation of astroglial mGluR3 promotes the non-amyloidogenic cleavage of APP by increasing the expression of ADAM10 and ADAM17 (α-secretases) and by decreasing BACE1 expression (β-secretase)^46^. We can then hypothesize that this disruption of mGluR3-APP complex could be related to an α-processing of APP after activation of mGluR3 as described in astrocytes (**Figure 8B**), rather than a conformational change of mGluR3.

To validate the amyloidogenic role of mGluR2, we chronically administered the DN13-DN1 nanobody, exhibiting PAM effects, to WT and 5xFAD mice. DN13-DN1 is selective for the mGluR2 homodimers and can only enhance its activation in the presence of glutamate, avoiding over-stimulation of the receptor. Treatment of mice with DN13-DN1 lasted 22 weeks (**Figure 5A**), time corresponding to the appearance of the cognitive deficits in 5xFAD male mice^47^. Behavioral tests show that the chronic positive modulation of mGluR2 worsen the spatial and short- and long-term memories in 5xFAD mice, while no impact was reported for WT mice (**Figure 5B-D**). The animal behavioral studies performed mainly target hippocampal neuronal functions. The chronic modulation with the DN13-DN1 nanobody exacerbate cognitive deficits in animals suggesting impairments of hippocampus. Indeed, we showed that deficits were associated with an increased number and surface of amyloid plaques in the hippocampus 5xFAD mice (**Figure 6A-F**). A minor effect of the chronic modulation of mGluR2 towards amyloid plaques in the cortex of 5xFAD mice, was observed, likely due to the decreased expression of mGluR2 homodimers only seen in the cortex (**Figure 7C**). The measurement of mGluR2 expression has some limitations as only the receptors at the plasma membrane are detected with the DN1 nanobodies. Thus, this mGluR2 decreased expression could be due to: (i) a decreased production of mGluR2 due to the disease progression; (ii) a modified internalization process of mGluR2; and/or (iii) an increased neuronal death. In addition, mice treated with DN13-DN1 exhibited lower levels of Aβ_1-42_ peptides in the soluble fractions (**Figure 6G**) associated with higher levels measured in insoluble fractions. Interestingly, treatment with DN1 nanobody, also decreased soluble Aβ_1-42_ peptides levels but these are not correlated with an increase in insoluble fraction. The chronic administration of DN13-DN1 to WT mice did not increase the production of mouse Aβ_1-42_ (**Figure 6K**) suggesting that treatment with a PAM mGluR2 in other brain diseases, like schizophrenia, will not induce any Alzheimer’s-like phenotype linked to Aβ_1-42_ production and aggregation as observed in the present study. This is the first proof-of-concept that chronic modulation of mGluRs can regulate the amyloidogenesis in AD by modulating the production of Aβ_1-42_ peptides. Modulation of the production, rather than increasing the clearance of Aβ peptides, may limit the ARIA detected in the classical immunotherapies. According to our work, development of a NAM mGluR2 and a PAM mGluR3 nanobodies could be promising tools for the treatment of AD.

To our knowledge, this study is the first to perform a chronic treatment using allosteric nanobodies that aims to evaluate the impact on a neurological disorder. Nanobodies are hydrophilic which limits the eventual side effects due to off-target binding in comparison with the hydrophobic small molecule PAM^48^. We also performed a toxicological study on 27 different organs from WT mice chronically administered with either 10 mg/kg of DN13-DN1, 10 mg/kg of DN1 or with vehicle. No adverse effects were detected in this toxicological study suggesting the relative innocuity of repeated administration of nanobodies in WT animals.

Altogether, our work deciphered the clear contribution of mGluR2 versus mGluR3 in the production of Aβ_1-42_ peptides in the physiopathology of AD, suggesting that inhibition of mGluR2 could be a promising pharmacological strategy for the treatment of AD. In addition, we provided evidence that a chronic nanobody-based immunotherapy modulating mGluRs activity can be considered for future treatments for AD, but also for other brain diseases.

## Material and methods

### Reagents

LY379268 and LY341495 (Bio-techne Tocris), pertussis toxin (Merck-Millipore), lipofectamine 2000 (Invitrogen), Triton X100, thioflavin T and guanidine hydrochloride (Sigma-Aldrich), Snap-Lumi4-Tb, Halo-Red, Halo-Lumi4-Tb, d2-NHS and Lumi4-Tb-NHS (Revvity) were used in the present study.

### Production and purification of nanobodies

Plasmid constructs of the DN1 nanobody acting as a silent allosteric ligand (SAL) and of the bivalent biparatopic nanobody DN13-DN1 with a positive allosteric modulation (PAM) property, were already published^22,23^. Both nanobodies are specific to mGluR2 and their large scale production was previously described^22,23^. Briefly, plasmids encoding the DN1 and DN13-DN1 nanobodies were transformed in E. coli BL21DE3 strain (Life technologies). Then, 10 mL of the preculture was added in 1 L of LB and was incubated until reaching an OD_600_ of 0.6-0.7. The nanobody expression was induced with 1 mM Isopropyl β-D-1-thiogalactopyranoside (IPTG). Bacteria were collected, resuspended in ice-cold TES buffer to induce the periplasmic lysis. The His-tagged nanobodies were purified by using Ni-NTA purification column (Qiagen) followed by desalting step using disposable PD-10 desalting columns (Cytiva) and endotoxins were removed using Proteus NoEndoTM S (Generon).

### Production, secretion, and Aβ_1-42_ peptides assayed by TR-FRET in HEK293 cells

Human Embryonic Kidney 293 (HEK293) T type cells were used to measure the production and secretion of Aβ_1-42_ peptides upon activation of mGluR2 or mGluR3 receptors. Cells were grown in Dulbecco’s Modified Eagle’s Medium (DMEM, Life technologies) supplemented with 10% heat-inactivated fetal bovine serum (FBS, v/v, Sigma-Aldrich) and maintained in humidified atmosphere containing 5% CO_2_ at 37°C. Absence of mycoplasma was assessed every month using the MycoAlert mycoplasma detection kit (LT07-318, Lonza). HEK293 cells were transfected by electroporation with plasmids coding for either Snap-tagged human mGlu2 receptors (^ST^mGluR2) or human mGluR3 (^ST^mGluR3) alone, or co-transfected with the Halo-tagged human APP_695_ construct (^HT^APP). In all conditions, HEK293 cells were co-transfected with the high affinity glutamate transporter EAAC1. In each condition, 20 × 10^6^ electroporated cells were transferred into a 6-well plate (Corning, Fisher Scientific) coated with polyornithine, 3×10^6^ cells per well. Mock cells were transfected with empty plasmids. Twenty-four hours after transfection, each well was incubated with DMEM-Glutamax at 37°C for 1 h, then medium was removed and cells were incubated with either PBS; agonist LY379268 (5 µM) alone, or with a pre-treatment with antagonist LY341495 (10 µM), or with pertussis toxin (PTX, 0.2 µg/mL); DN13-DN1 nanobody (200 nM); or with the EC_20_ of LY379268 (0.5 nM) alone, or combined with DN13-DN1. After 12 h of treatment, the cell supernatant was collected and centrifuged 10 min at 1,000 g to remove cell debris. Supernatants were supplemented with a protease inhibitor cocktail (Complete ultra, Roche, Sigma-Aldrich) and stored at −80°C, until use. The human Aβ_1-42_ peptides was assayed with a FRET-based assay (Revvity). FRET signal was determined by measuring the sensitized acceptor emission (665 nm) and Europium donor emission (620 nm) using a 50 μs delay and 450 μs integration time upon excitation at 337 nm on a Pherastar FS plate-reader (BMG LabTech). Then, acceptor/donor ratio (665 nm/620 nm × 10^4^) was calculated and converted into pg/mL values according to the standard curve of the assay. All values obtained were normalized by the Mock condition and expressed as a fold of basal.

### Cell surface expression of transfected mGluR2, mGluR3 and APP in HEK293 cells

Part of the HEK293 cells transfected described above were also transferred in a 96-well plate (Greiner bio-one, PS, F-bottom), 10^5^ cells per well. Twenty-four hours after transfection, cells were incubated with 100 nM of Snap- or Halo-Lumi4-Tb substrates in DMEM-Glutamax for 1 h at 37°C, to reveal mGluRs or APP (referred as t=0 h). Another set of cells was washed with DMEM-Glutamax for 1h at 37°C, incubated with the different drugs cited above for 12 h at 37°C, and then incubated with the appropriate Lumi4-Tb (referred as t=12 h). The cell surface relative expression of the different receptors was measured using the Terbium donor emission at 620 nm, upon excitation at 337 nm on a Pherastar FS plate-reader.

### TR-FRET saturation assay

FRET saturation assays were performed as previously described^27^ to determine the potential association of APP with mGluR2 or mGluR3. Briefly, HEK293 cells were co-transfected by lipofection (Lipofectamine 2000) with a constant amount of plasmid encoding ^ST^APP, ^ST^mGluR2, ^ST^GB1a-ASA or ^ST^mGluR3 (20 ng), and increasing amounts of ^HT^mGluR2, ^HT^mGluR3, ^HT^mGluR5, ^HT^APP, ^HT^mGluR3, ^HT^mGluR3(Del829-879), ^HT^mGluR3(Del570-879), ^HT^APP(Del648-695), ^HT^mGluR2-R3 Cterm or ^HT^mGluR3-R2 Cterm (from 0 to 160 ng). Cells were plated in 96-well plates coated with polyornithine at 10^5^ cells per well. Twenty-four hours after transfection, cells were incubated with DMEM-Glutamax 1 h at 37°C containing 100 nM of Snap-Lumi4-Tb to measure ^ST^receptors expression (referred as Exp ST), Halo-Lumi4-Tb to measure ^HT^receptors (Exp HT), or with a combination of 100 nM of Snap-Lumi4-Tb and 100 nM of Halo-red to measure the FRET signal. Cells were then washed with Tag-light buffer and fluorescence as well as TR-FRET were read using an Infinite F500 spectrofluorometer (Tecan).

### Diffusion-enhanced resonance energy transfer (DERET) internalization assay

A DERET assay^28^ was performed to measure the internalization in real time of the human APP upon activation of mGluR2 or mGluR3 (Sup Figure 2). The internalization assay was performed and adapted as previously described^21,49^. HEK293 cells were transfected by lipofection either with ^ST^mGluR2, ^ST^mGluR3 or ^HT^APP alone, or co-transfected with ^ST^mGluR2 or ^ST^mGluR3 with the ^HT^APP. In all conditions, the high affinity glutamate transporter EAAC1 was also transfected. Cells were transferred in black non-transparent 96-well plates at 10^5^ cells per well. Twenty-four hours after transfection, receptors were labeled with either 100 nM of Snap-Lumi4-Tb or Halo-Lumi4-Tb in Tag-lite buffer, for 1.5 h at 4 °C. Excess of Snap- or Halo-Lumi4-Tb substrates was removed by washing cells with Tag-lite buffer. The internalization assay was performed by incubating the cells with the Tag-lite buffer, either with PBS or with 1 µM of LY376892 and in the presence of an excess of fluorescein (24 µM). Lumi4-Tb was excited at 337 nm, and the emission fluorescence intensities were recorded for the donor (620 nm, 1500 μs delay, 1500 μs reading time) and acceptor (520 nm, 150 μs delay, 400 μs reading time) using a pherastar FS microplate reader. Both intensities (620 nm and 520 nm) were measured every 5 min for 1.5 h. The ratio of 620 nm/520 nm was then calculated for each time point and multiplied by 10^4^.

### Ethics

This project follows the specific French national guidelines on animal experimentation and well-being according to the European Directive 2010/63/EU. This project was approved by the French National Ethic Committee for Animal Experimentation and the French Ministry of Agriculture under the number APAFIS #28773-2020121811289735v4. Animals were housed under a 12 h light/12 h dark cycle and at 23 ± 2 °C. Animals had free access to water and food and were fed under a standard chow diet (A03) (SAFE Diets).

### Animals

5xFAD mice overexpress the human APP with the Swedish (K670N, M671L), Florida (I716V) and London (V717I) mutations and the human Presenilin1 (PS1) gene harboring the M146L and L286V mutations^47^. These transgenes are regulated by the neuronal-specific elements of the mouse Thy1 promoter. 5xFAD heterozygous transgenic mice were used for the experiments and WT littermates as controls. All animals were genotyped by qPCR using tail genomic DNA^47^. To study the effect of a chronic positive modulation of mGluR2 *in vivo*, a nanobody specific of mGlu2 receptors and exhibiting PAM effects^23^ (DN13-DN1) was used. As a negative control, we used the DN1 nanobody^22^, with a neutral effect on mGluR2 activation. In this study, 4-weeks-old male mice were chronically treated with 10 mg/kg of either DN13-DN1 (n=20 WT and n=21 5xFAD) or DN1 (n=18 WT and n=16 5xFAD), by intraperitoneal route, once a week for 22 weeks. Control mice were treated with an equivalent volume of PBS (n=16 WT and n=22 5xFAD). Mice were followed up and weighted every week. After treatment, mice underwent several behavioral tests and were sacrificed.

## Behavioral tests

One week before behavioral tests, mice were handled every day by the operator. Mice groups were as such: DN13-DN1 nanobody treatment (n=17 WT and n=17 5xFAD), DN1 treatment (n=15 WT and n=11 5xFAD) and PBS (n=16 WT and n=22 5xFAD).

### Nest building

Animals, housed in group, were placed in individual testing cages and received 3 g of nestlet for an overnight nest building. The next morning before 9:00 a.m., nest quality was scored, and non-shredded nestlet pieces were weighed. Scoring of the nest quality was performed using a 5-point nest-ratting scale^29^.

### Marble-burying

A plastic cage (width: 23.4 cm, length: 37.3 cm, height: 14.0 cm) was filled with 5 cm of bedding. 18 marbles were placed on top of this bedding, 3 marbles in 6 rows. Each mouse was placed in an individual testing cage and was left with the marbles for 30 min. Then, mouse was removed, and the number of buried marbles was counted. We considered a marble buried when at least two-thirds of the marble was covered of bedding as previously described^50^.

### T-maze

The T-maze is composed of one main runway (width: 10 cm, length: 50 cm, height: 14.0 cm) connected to two side arms (width: 10 cm, length: 20 cm, height: 14.0 cm) made of acrylic plastic painted in light grey color. During the T-maze spatial habituation, each mouse was allowed to explore for 5 min the main runway and one of the side arms. Mice were placed back to their home cages for 5 min. Then, the second (novel) side arm was opened and the ambulation of mice in each side arm was observed for 5 min. The number of entries in each side arms was quantified, and the discrimination index was calculated as “(novel − familiar) / (familiar + novel)”.

### Novel object recognition

This behavioral test was performed as previously described^51^. White acrylic boxes (width: 28 cm, length: 41 cm, height: 28 cm) and three distinct objects with different shapes were used. During all the behavioral period, the room brightness was dimmed (30 lux) and bedding was spread on the floor of the apparatus, to reduce anxiety. Each day, prior to begin the behavioral test, mice were familiarized with the testing room, for at least 1 h. On the first day, all mice were exposed to the empty arena for 5 min for habituation. The next day, during the familiarization phase, all mice were exposed to two identical objects for 5 min. Four hours after the familiarization, one object was kept as a familiar object, while the other was replaced by a novel object. Mice were allowed to explore the objects for 5 min. The same procedure was performed 24 h after familiarization by using a different novel object than the one used at 4 h.

For the habituation trials, the mean velocity, the time spent in the center of the arena, the total distance traveled, and the time of immobility were measured using EthoTrack software (v. 3.11.0., Innovation Net). For the other trials, a discrimination index using the exploration times ([novel − familiar] / [familiar + novel]) was calculated. Direct contact or sniffing behavior close to objects (< 1 cm) were defined as object exploration, while climbing and digging were not considered. Mice showing a preference for one of the familiar objects or having a total exploration time of the objects of less than 6 s during trials, were excluded from behavioral analyses. Two independent quantifications of the discrimination indexes were performed blindly, and the same trends were observed for the two sets of values obtained. Results from the two independent quantifications were averaged for each mouse.

### Culling of mice and tissue collection

This procedure was performed as previously described^52^. Briefly, mice were anesthetized (16% ketamine, 4% xylazine in sodium chloride 0.9%) and sacrificed by an intracardial perfusion of PBS. Brains were removed and cut along the sagittal axis. Left hemispheres were directly frozen in liquid nitrogen and stored at −80°C (for biochemical analyses), and right hemispheres were in part fixed in a 4% paraformaldehyde solution (PFA, Euromedex) overnight at 4°C (for immunolabeling). Fixed hemibrains were rinsed with PBS and incubated in a 30% sucrose solution for 4 days at 4°C. Brains were then included in O.C.T. (Tissue-Tek®, Sakura Finetek), quickly frozen on acetone chilled on dry ice, and conserved at −80°C.

The other part of the right hemispheres was collected to perform TR-FRET experiments as described in Meng et al^20^. For each hemibrain, different regions were punched: cortex, hippocampus, cerebellum, and the midbrain. Samples were collected in a 1.5 mL cryogenic tube (ThermoFisher Scientific) in 1 mL cold cryopreservation medium (DMEM-Glutamax supplemented with 10% FBS and 10% DMSO), frozen at −80°C in a freezing box and kept at −80 °C until use.

In parallel, in 2 WT mice from each group (PBS, DN13-DN1 and DN1), 27 organs were collected (eyes with optical nerve, harderian glands, salivary glands, stomach, pancreas, intestines (duodenum, ileum, jejunum, and colon), cecum, testes, epididymis, seminal vesicles, prostate, urinary bladder, kidneys, adrenal glands, white adipose tissue, spleen, liver, heart, aorta, lungs, esophagus, trachea, bone with bone marrow, lymph nodes, sciatic nerve, skeletal muscle, and skin). All the samples were incubated overnight in a 4% PFA solution at 4°C, washed with PBS and kept in 70 % ethanol. Samples were processed for anatomopathological studies according to the international RITA procedure for toxicological studies^53–55^ and were dehydrated in graded ethanol and embedded in paraffin.

### Brain extracts preparation

Frozen left hemispheres of 5xFAD (22 vehicle, 17 DN13-DN1 and 11 DN1) were thawed, weighed, and homogenized with a potter in 20% wt/vol in a tris-saline buffer (20 mM Tris-HCl, pH 7.4; 150 mM NaCl) containing a protease inhibitor cocktail (Complete ultra). Homogenates were then ultracentrifuged at 355,000 g at 4°C for 20 min (TLA 120.1 rotor, Beckman Optima TLX ultracentrifuge) and supernatants (referred as “soluble fraction”) were collected and kept at −80°C, until use. Pellets were resuspended by brief sonication in 6 M guanidine HCl and 20 mM Tris-HCl (pH 7.6) buffer and ultracentrifuged at 227,000 g at 4°C for 20 min. Supernatants obtained (referred as “insoluble fraction”) were collected and stored at −80°C. For WT mice hemibrains (12 vehicle, 12 DN13-DN1 and 10 DN1), the same procedure was performed.

### Amyloid-β assays

FRET-based assays for the dosage of human Aβ_1-40_ (#62B40PEG) and Aβ_1-42_ (#62B42PEG) were used according to the manufacturer’s instructions (Revvity). FRET signal was determined as explained above. Mouse Aβ_1-42_ were assayed using an Elisa kit from ThermoFisher Scientific (#KMB3441). Reactions were read at 450 nm using an Infinite M200 (Tecan). All values were normalized according to their protein concentration determined by the bicinchoninic acid (BCA) method protein assay (Sigma-Aldrich). BCA OD at 562 nm was measured using an Infinite M200 (Tecan).

### Aggregate staining and quantification

Frozen hemibrains included in OCT were mounted on a cryostat (Leica) and 20 μm coronal sections were performed and conserved at −20°C in an antifreeze solution. Brain tissue sections were incubated with a Thioflavin T at 0.01% for 10 min^52^. Sections were washed with an 80% ethanol solution, rinsed with distilled water and mounted using Fluoroshield containing DAPI to stain nuclei (Sigma-Aldrich). All sections were visualized using a slide scanner Axio scan Z1 microscope (Zeiss) by performing full-section mosaic with a 20x enlargement. Images were analyzed with Fiji software (version 2.0; National Institutes of Health) and the number and the surface of amyloid aggregates were quantified as previously described^52^. The number of brains used per group to quantify the amyloid plaques is the following: DN13-DN1 (n=6 WT and n=7 5xFAD), DN1 (n=7 WT and n=6 5xFAD) and vehicle (n=8 WT and n=8 5xFAD).

### HES staining for toxicological study

The 27 samples collected from WT mice and embedded in paraffin were cut using a microtome and 5 μm tissue sections were performed. All sections were mounted on Superfrost Plus slides, were dewaxed and stained with hematoxylin (nucleus staining), eosin (cytoplasm staining) and natural saffron (collagen fibers staining) (HES). Sections were washed, dehydrated and then coverslipped with Permount mounting medium (Fisher Scientific). All sections were visualized using a slide scanner Axio scan Z1 microscope by performing full-section mosaic with a 20x enlargement.

### Nanobody coupling to fluorophores

The mGluR2 specific nanobody DN1 was labelled as previously described^20^. Briefly, the nanobody was dialyzed overnight at 4°C and 250 μg of nanobody were incubated at 20°C with d2-NHS (Acceptor) in 0.1 M carbonate buffer (pH = 9) at a molar ratio of 6, for 45 min. In parallel, 250 µg of the dialyzed nanobody were incubated with Lumi4-Tb-NHS (donor) in 50 mM phosphate buffer (pH 8) at a molar ratio of 12 for 30 min. Coupled nanobodies were then purified using a gel filtration column in 100 mM phosphate buffer at pH 7 and their concentration was determined by OD at 280 nm. After purification, the labelled nanobodies were supplemented with 0.1 % BSA and stored at −20°C, until use.

### Relative quantification of the mGluR2 homodimers by TR-FRET in brains from WT and 5xFAD mice

For this experiment, the number of hemibrains analyzed was as follows: DN13-DN1 (n=6 WT and n=8 5xFAD), DN1 (n=3 WT and n=3 5xFAD) and PBS (n=5 WT and n=10 5xFAD). The TR-FRET protocol was performed as described in Meng et al ^20^. Samples were rapidly thawed in a water bath at 37°C and tissues were washed 2 times with DMEM Glutamax and 2 times with cold PBS. Tissues were then digested with 200 μL Versene solution (ThermoFisher) for 20 min at 20°C, and the cells were dissociated by several pipetting. Several rounds of cell dissociation in DMEM Glutamax were performed to get the maximum number of cells. Then, dissociated cells were centrifuged at 3,000 g for 5 min, and the cell pellet was washed with 1 mL of cold PBS and resuspended in 200 μL cold PBS. Then, 10 μL of cells were added in low volume 96-well microplate (Revvity) and incubated with 10 µL of DN1-d2 (25 nM), 10 µL of DN1-Tb (25 nM) and 10 µL of LY341495 (10 µM), to reach a total volume of 40 μL. Plates were incubated overnight at 22°C and the TR-FRET signal was determined by measuring the sensitized acceptor emission (665 nm) and donor emission (620 nm) using a 50 μs delay and 450 μs integration time upon excitation at 337 nm on a Pherastar FS plate-reader. In parallel, the total amount of proteins was determined by the BCA protein assay (Sigma-Aldrich). Absorbances were measured using an Infinite M200.

### Statistical analyses

All data were initially analyzed on GraphPad Prism software (version 10.0; GraphPad) using Shapiro-Wilk’s normality test. When data were normally distributed (*p* > 0.05), statistical analyses were performed using parametric tests: one-way ANOVA (Holm-Sidak’s *post-hoc* test) or the two-way ANOVA (Holm-Sidak’s *post-hoc* analysis). When data were not normally distributed (*p* < 0.05) statistical analyses were performed using a nonparametric test: Kruskal-Wallis (Dunn’s *post-hoc* test). A probability of 0.05 has been defined as a significant difference for all statistical analyses.

## Acknowledgements

We thank Meng Jyiong for technical advice for the relative quantification of mGluR2 in brains, Cécile Derieux for her experimental support and advice, Dr. Nigel Moore for the tissue histopathological analysis and Revvity for the reagents. We acknowledge the RHEM facility (Réseau d’Histologie Expérimentale de Montpellier, IRCM, Montpellier) for processing samples, histology staining and expertise for the RITA procedure; ARPEGE facility (Pharmacology Screening-Interactome, Institut de Génomique Fonctionnelle, Montpellier) for their assistance in measurements; the imaging facility MRI (Institut des Neurosciences de Montpellier) for their help in image acquisition and analysis; and the animal facility RAM-iExplore from BioCampus (Montpellier, France) and Michel Salvado for technical assistance with animals. V.P. and P.R. were supported by the Centre National de la Recherche Scientifique (CNRS), the Institut National de la Santé et de la Recherche Médicale (INSERM), and by grants from the Agence Nationale de la Recherche (ANR-22-CE18-0003), and the FRM (FRM team: DEQ20170336747). P.-A.L. was the recipient of a postdoctoral fellowship from the LabEX MAbImprove (grant NR-10-LABX-5301); C.E.P. was the recipient of a fellowship from the French Ministry of Higher Education and Research; M.E.T. was supported by a doctoral fellowship from the LabEX MAbImprove (grant NR-10-LABX-5301) and from the Center of Excellence for Neurodegenerative diseases of Montpellier (CoEN) – CHU of Montpellier.

## Author contributions

P.-A.L., J.-P.P., V.P., J.L., L.P. and P.R. designed experiments; S.C. contributed to the design of mouse experiments; P.-A.L. performed Aβ assay in cellular experiments; P.-A.L. performed FRET saturation assays; P.-A.L. performed DERET internalization assays; P.-A.L., J.M., S.R., M.E.T., S.D., P.C., M.O., J.K. and U.A. performed nanobodies production; P.-A.L., J.M., S.R., S.D. and U.A. performed genotyping of mice, injection of mice, culling of mice and tissue collection; P.-A.L. performed behavioral tests; P.-A.L., J.B. and S.D. performed cryosectioning of brains; P.-A.L. and J.B. performed histology and brain Aβ assays; P.-A.L., J.B., M.E.T., S.D. and U.A. performed relative quantification of mGluR2; P.-A.L., C.E.P., M.C. and J.B. performed data analysis; P.-A.L., V.P. and P.R. wrote the manuscript. All authors reviewed the manuscript.

## Competing interests

P.R. and J.-P.P. are involved in a collaborative team between the CNRS (IGF, Montpellier) and Revvity. The remaining authors declare no competing interests.

## Notes

### Competing Interest Statement

P.R. and J.-P.P. are involved in a collaborative team between the CNRS and Revvity Cisbio. The remaining authors declare no competing interests.

