## Supplementary figures 1-6 for "Chronic potentiation of metabotropic glutamate receptor 2 with a nanobody accelerates amyloidogenesis in Alzheimer’s disease"

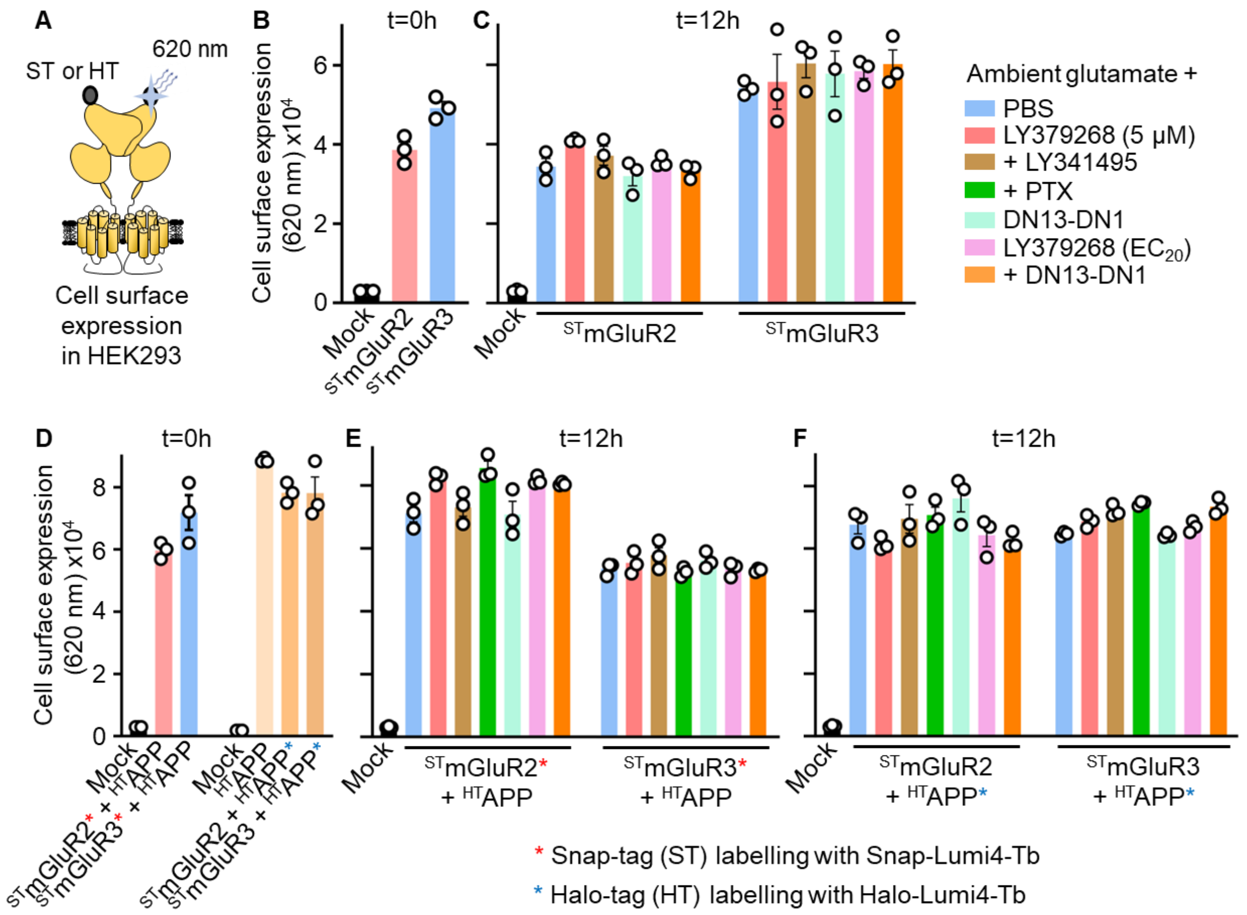


**Supplementary Figure 1. Cell surface expression of transfected receptors. A.** The expression of transfected $^{ST}$mGluR2 and $^{ST}$mGluR3 at the surface of HEK293 cells was measured using Snap-Lumi4-Terbium (Tb) and for $^{HT}$APP by using Halo-Lumi4-Tb. Expression was measured using the Tb emission at 620 nm, upon excitation at 337 nm. **B-F.** Expression of $^{ST}$mGluR2, $^{ST}$mGluR3 or $^{HT}$APP prior to any treatment of cells (t=0 h, **B, D**) or after 12 h of treatment with the different compounds as described in Figure 1 (**C, E-F**). Expression levels in **B-C** are related to Figure 1B, and expression levels in **D-F** are relative to Figure 1C. Expression levels of mGluR receptors or APP is equivalent in all conditions. Data are presented as mean ± SEM from 3 biologically independent experiments performed in triplicates.

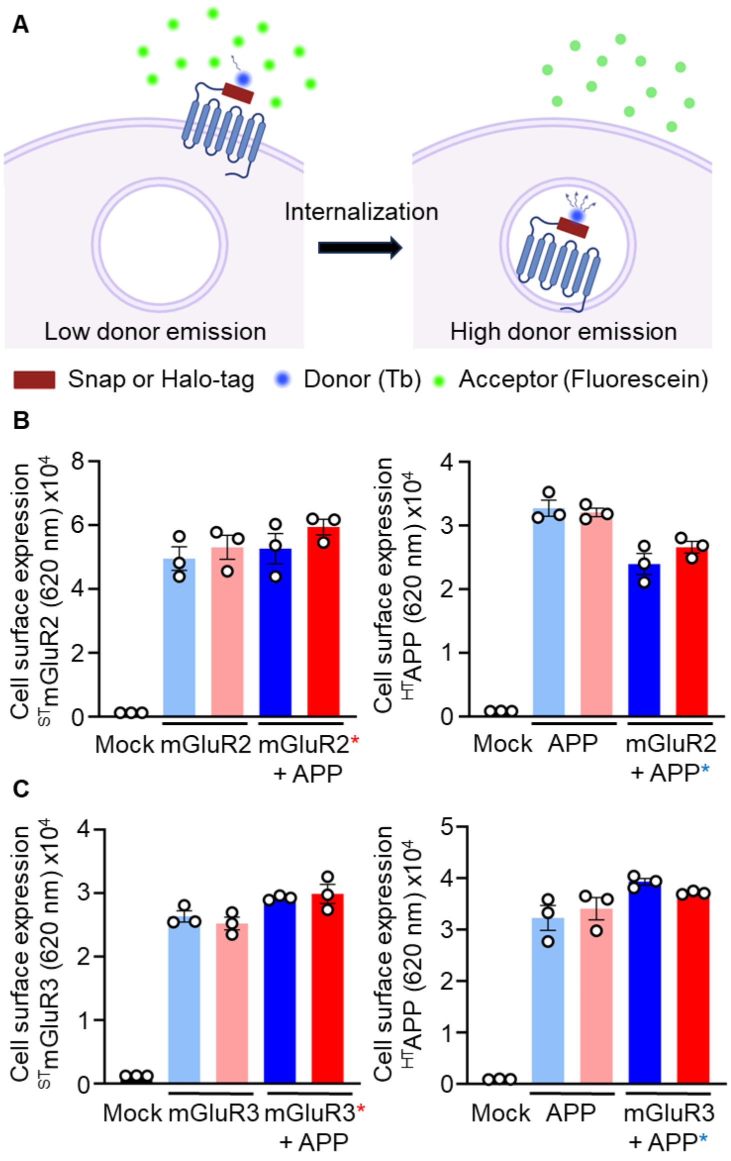


**Supplementary Figure 2. DERET internalization assay and cell surface expression of transfected receptors A.** Schema illustrating the DERET-based internalization assay. Created with BioRender.com. **B-C.** Cell surface expression of $^{ST}$mGluR2 and $^{HT}$APP (**B**) and $^{ST}$mGluR3 and $^{HT}$APP (**C**), either transfected alone or co-transfected in HEK293 cells. When co-transfected, mGluR receptors were labelled with Snap-Lumi4-Tb (*) whereas APP was labelled with Halo-Lumi4-Tb (*). Expression levels in **B** are related to Figure 4A, and expression in **C** to Figure 4B. Data are presented as mean ± SEM from 3 biologically independent experiments performed in triplicates.

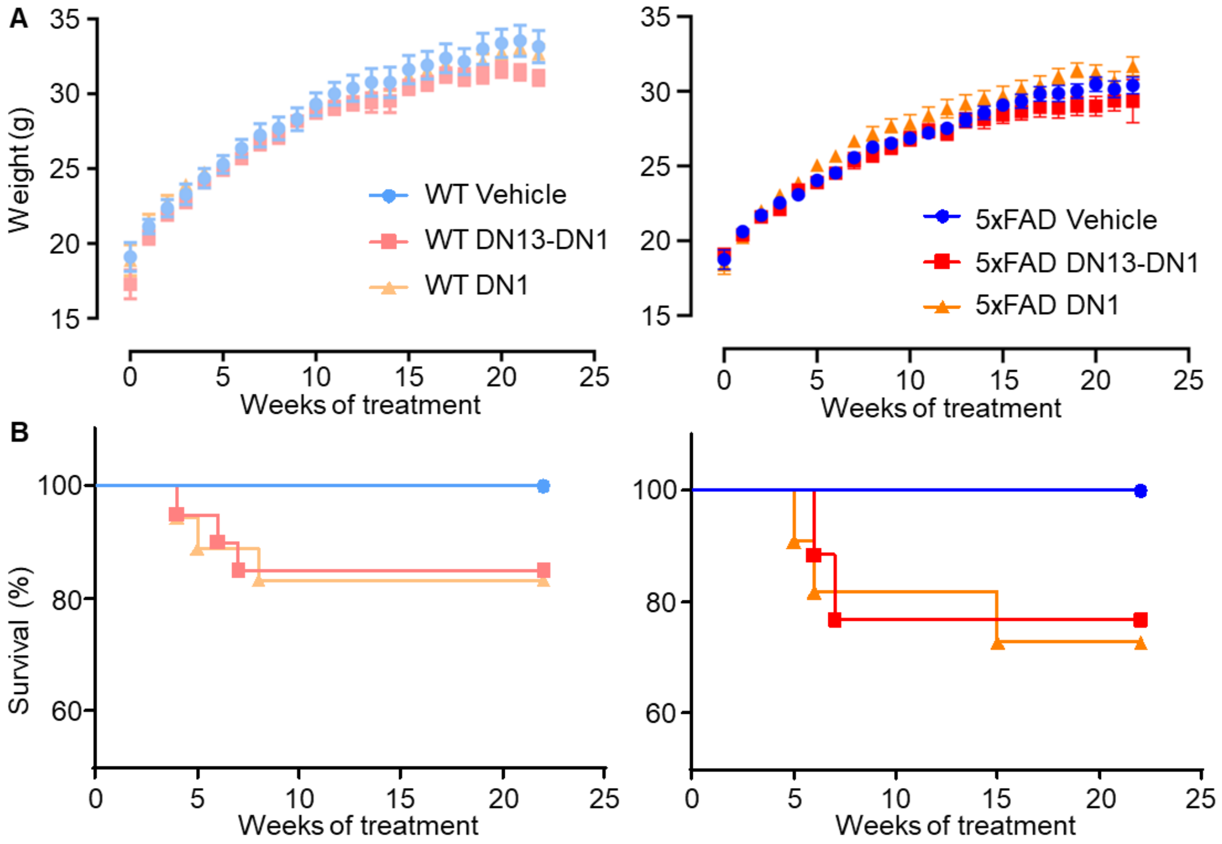


**Supplementary Figure 3. Weight and survival of WT and 5xFAD mice.** Weight evolution (**A**) and survival curves (**B**) of WT and 5xFAD mice treated with 10 mg/kg of DN13-DN1, 10 mg/kg of DN1, or with an equivalent volume of vehicle, for 22 weeks. Data are presented as mean ± SEM (**A**) or as a percentage of survival (**B**).

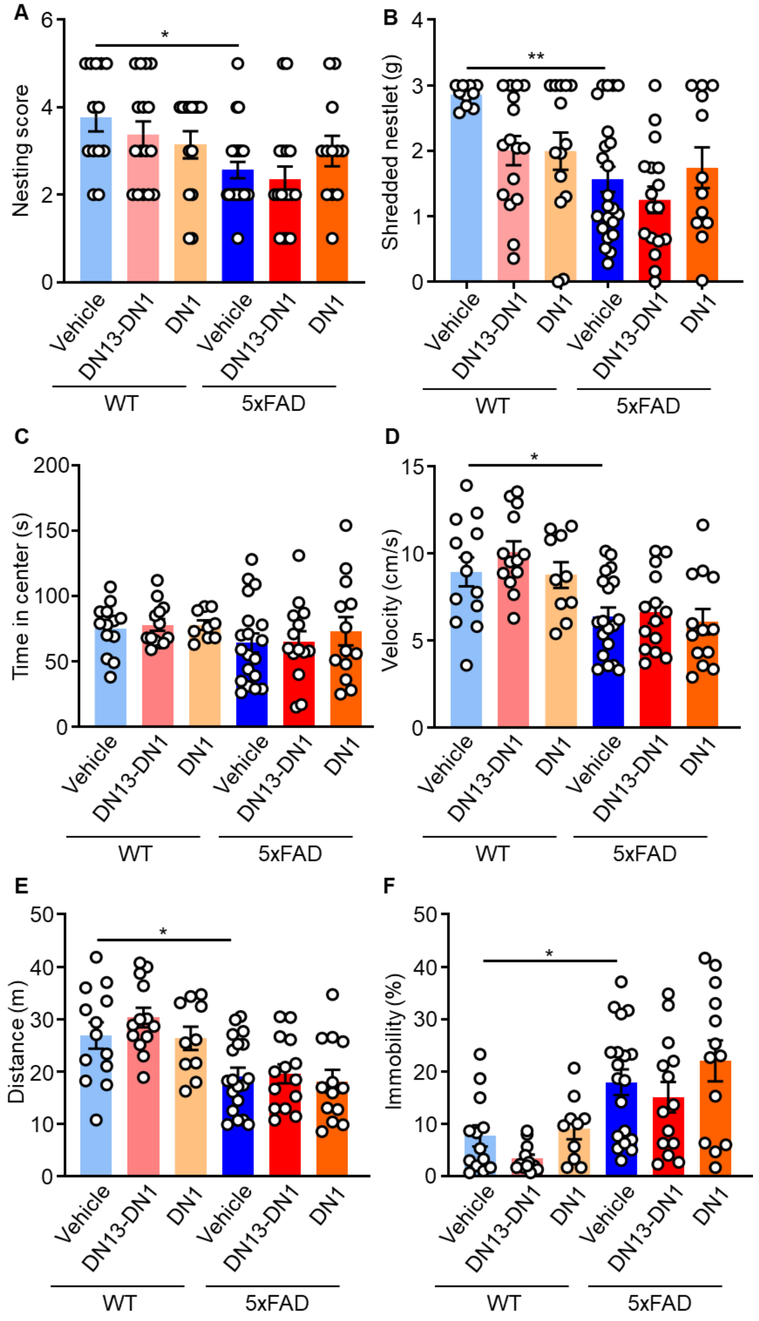


**Supplementary Figure 4. Treatment with nanobodies does not influence nesting and locomotor activity in 5xFAD and WT animals. A-B.** Nesting score (**A**) and quantity of shredded nestlet (**B**) in the nest building test for WT and 5xFAD mice treated with vehicle, 10 mg/kg of DN13-DN1 or with 10 mg/kg of DN1 nanobodies. **C-F.** Velocity (**C**), time spent in the center of the arena (**D**), total distance (**E**) and percentage of immobility

(**F**) in the arena during the habituation day of the novel object recognition test for WT and 5xFAD mice treated with vehicle, 10 mg/kg of DN13-DN1 or with 10 mg/kg of DN1 nanobodies. All data are presented as mean ± SEM (n = 12-21 mice/group) and analyzed using the two-way ANOVA followed by a Holm-Sidak's *post hoc* analysis (* $p < 0.05$, ** $p < 0.01$, *** $p < 0.001$).

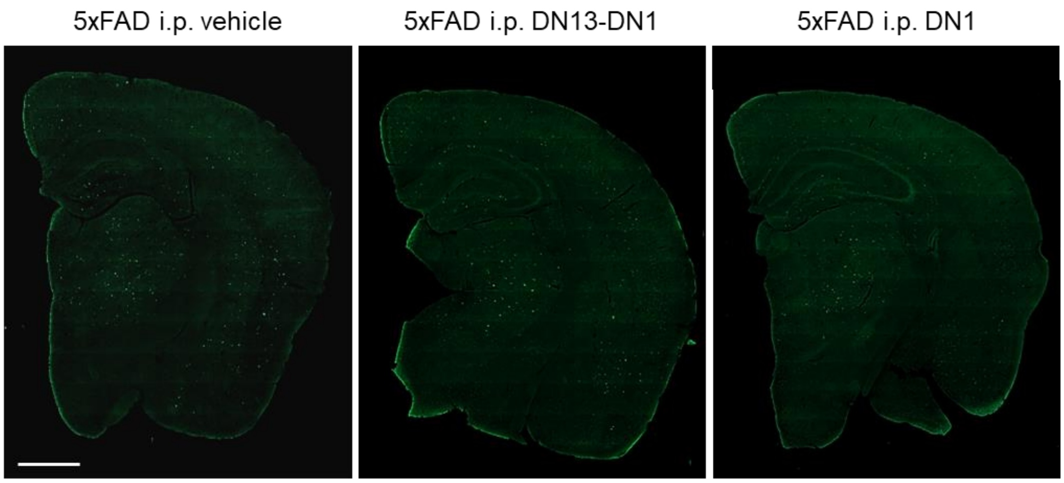


**Supplementary Figure 5.** Representative images of ThT positive amyloid plaques in the hemibrains of 5xFAD mice (27 weeks old) chronically treated with 10 mg/kg of DN13-DN1, 10 mg/kg of DN1 or with vehicle. Images are mosaics obtained using a slide scanner Axio scan Z1 microscope with a 20X objective (scale bar: 1 mm).

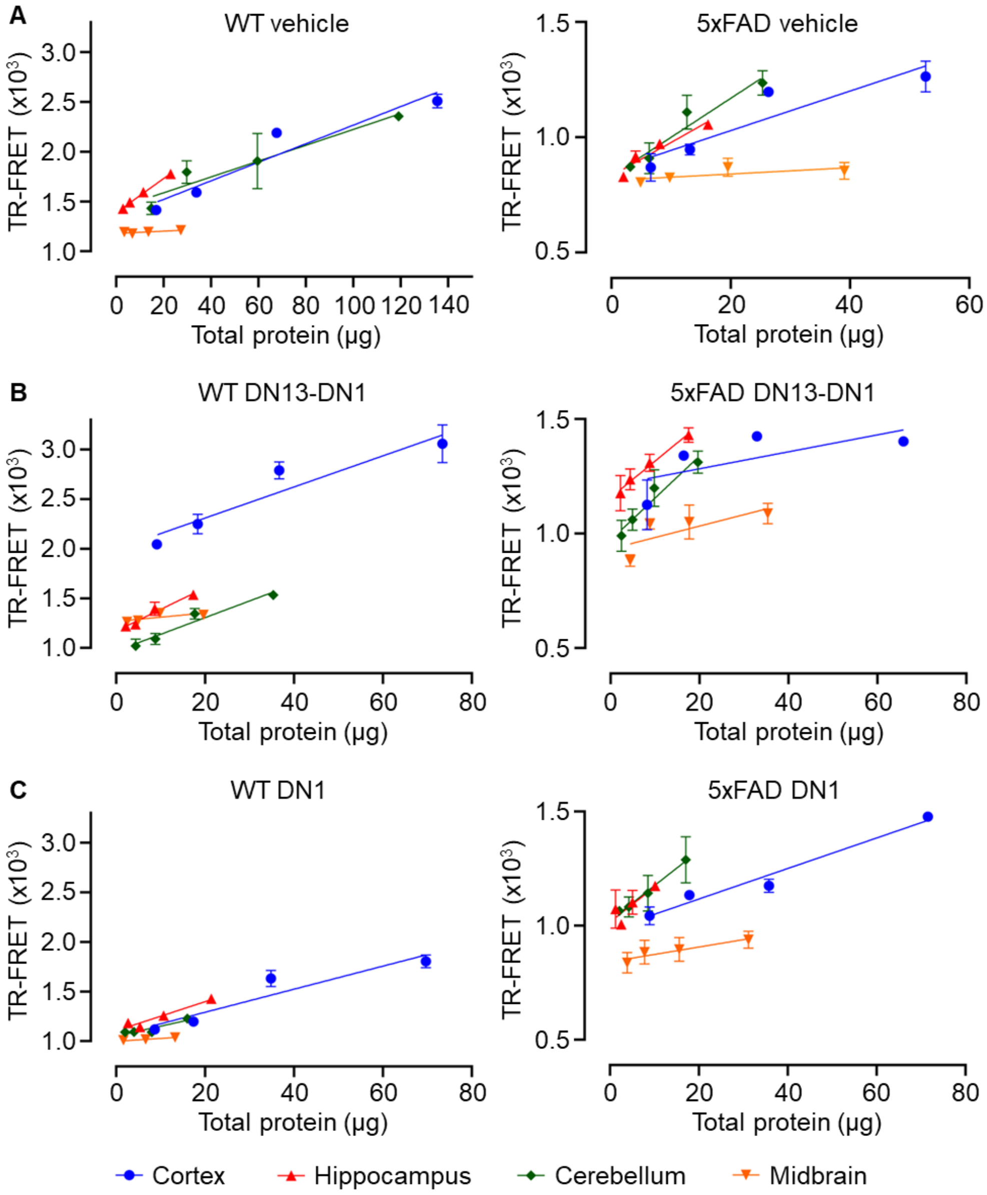


**Supplementary Figure 6.** Representative relative linear quantification experiments of mGluR2 homodimers presented in Figure 7 of WT and 5xFAD mice chronically injected with vehicle (**A**), with 10 mg/kg of either DN13-DN1 (**B**) or DN1 nanobody (**C**), for the different regions, cortex, hippocampus, cerebellum, and midbrain. Data are presented as mean ± SEM of a triplicate from one representative animal of each group of treatment.

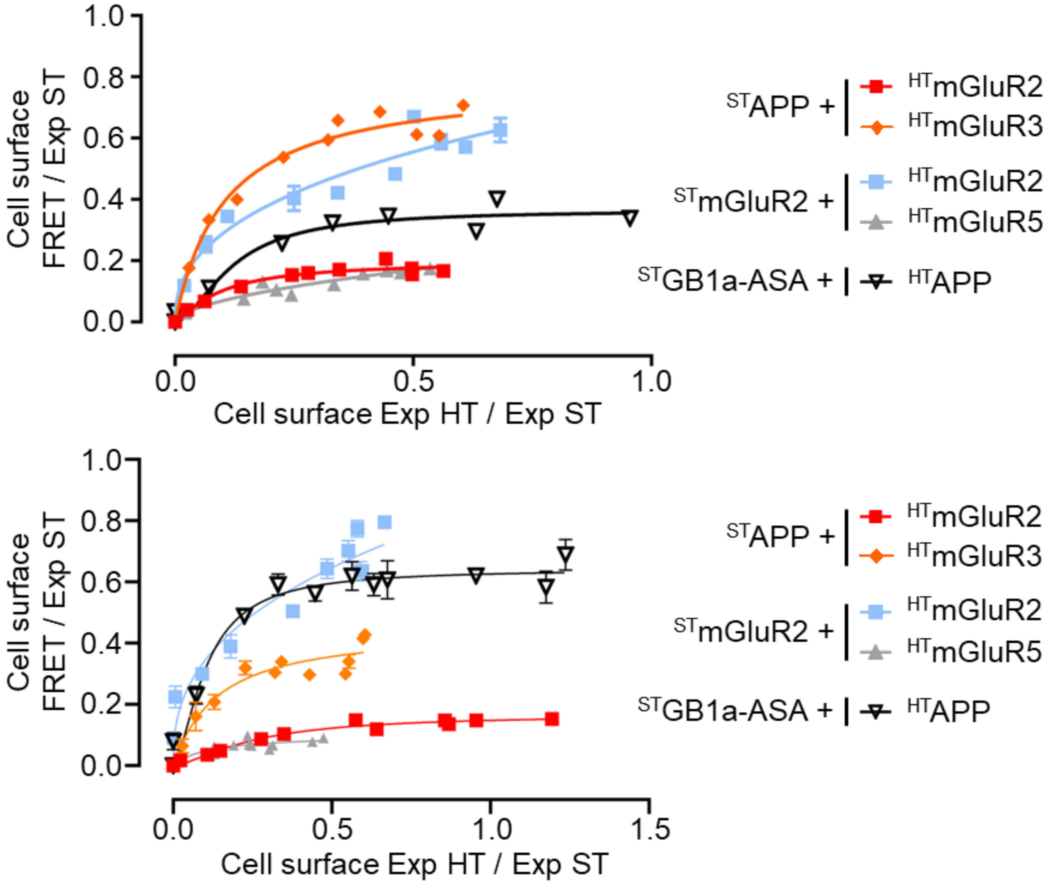


**Source data 1.** TR-FRET saturation curves of the two other experiments performed in Figure 2. Data are presented as mean ± SEM of triplicates. Pictograms for the different conditions, tags and labelling used are identical as in Figure 2.

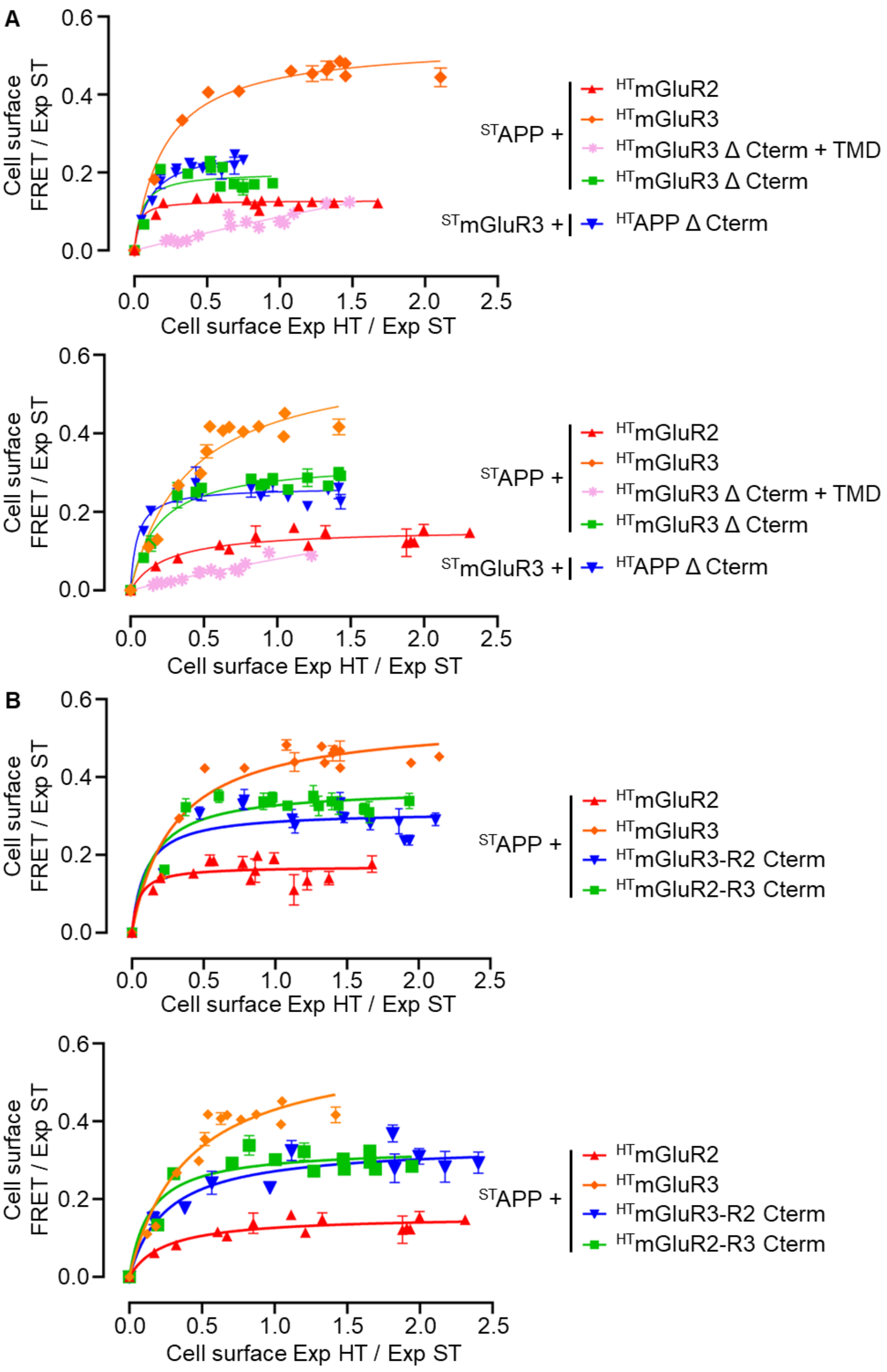


**Source data 2.** TR-FRET saturation curves of the two other experiments performed in Figure 3A (**A**) and Figure 3B (**B**). Data are presented as mean ± SEM of triplicates. Pictograms for the different conditions, tags and labelling used are identical as in Figure 3.
